# Biological polyethylene deconstruction initiated by oxidation from DyP peroxidases

**DOI:** 10.1101/2025.02.27.640435

**Authors:** Ross R. Klauer, D. Alex Hansen, Zoé O. G. Schyns, Lummy Oliveira Monteiro, Jenna A. Moore-Ott, Mekhi Williams, Megan Tarr, Jyoti Singh, Ashwin Mhadeshwar, LaShanda T. J. Korley, Kevin V. Solomon, Mark A. Blenner

## Abstract

Polyethylene (PE) is the most commonly used plastic on Earth due to its favorable material properties such as high ductility, mechanical strength, and bond homogeneity that make the material resistant to deconstruction. However, the lack of robust recycling infrastructure for PE end-of-life management is leading to an estimated 4 million tons of environmental accumulation annually, with implications for human and environmental health. Biological deconstruction and upcycling could potentially aid in PE waste management by allowing for high-yield conversion of waste plastics to high value products, although such processes are not yet possible. In this work, we mined the gut of low-density PE (LDPE) fed mealworms that can reduce LDPE molecular weight by >40% and discovered dye decolorizing peroxidases (DyPs) that oxidized LDPE, initiating biological deconstruction. A plastic-active DyP is characterized by a hydrophobic loop near its active site that helps mediate binding and tunes activity. LDPE oxidation is driven by surface exposed residues proximal to the active site enabling activity on polymeric substrates. Our work provides robust evidence for enzymatic LDPE deconstruction and identifies molecular targets for further development to realize scalable biological LDPE upcycling.

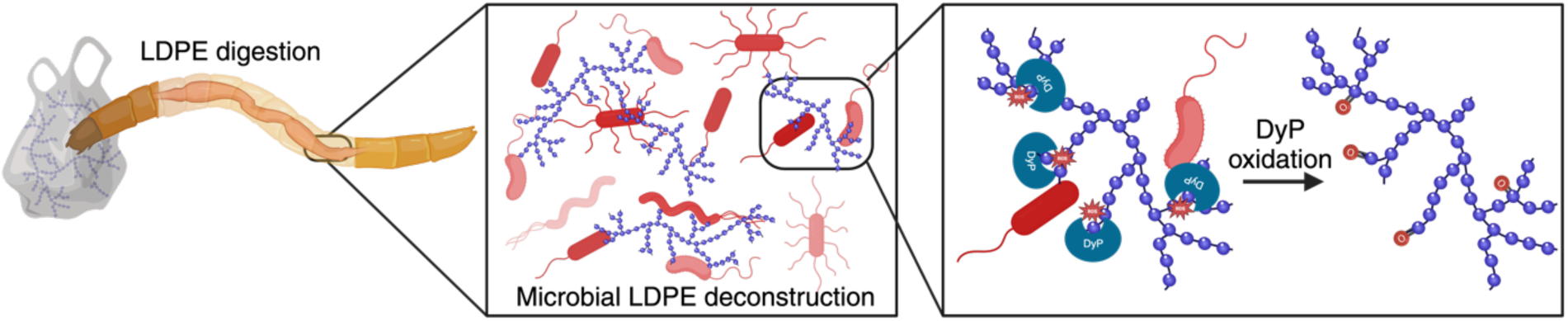

## Introduction

More than 300 million tons of plastics are produced globally each year of which >90% are not effectively recycled; this plastic waste ends up in the environment, landfilled, or incinerated, generating negative environmental impacts and increasing greenhouse gas emissions (*1–3*). Mechanical recycling can result in recycled plastics with inferior properties compared to virgin plastics and thus reduced value after many cycles (*3*, *4*). Chemical recycling and upcycling strategies produce virgin-quality recycled plastics through deconstruction and repolymerization or upcycled products through deconstruction and new synthesis (*3*, *5*). These technologies are promising but often require high temperatures and pressures for operation and use expensive catalysts that limit industrial adoption (*6*, *7*). Biological plastics waste valorization offers alternative approaches to chemical recycling with lower operating temperatures and pressures and still provides the opportunity to upcycle plastics waste into higher value products (*8*, *9*). Recent rapid progress in identifying and engineering poly(ethylene terephthalate) (PET) degrading enzymes have resulted in impressive rates of PET deconstruction up to 61.3 g_PET_L^-1^h^-1^ and the first commercial-scale PET biological recycling plant (*10–14*). Contrary to PET, methods for the biological deconstruction of non-hydrolysable plastics — which make up approximately 70% of plastics waste — remain undeveloped (*4*).

The most abundant plastics, polyethylene (PE) and polypropylene (PP) (*4*), are composed exclusively of C-C backbones that lack bonds susceptible to enzymatic hydrolysis by enzymes such as esterases or hydrolases, meaning that they are difficult to deconstruct biologically. Moreover, these C-C bonds are stronger than the labile ester or amide bonds found in hydrolysable plastics (*15*). Therefore, C-C bonds must be activated through functionalization such as oxidation to allow for their cleavage. While enzymes have been reported to act on non-hydrolysable plastics (*16*, *17*), these reports have been met with healthy skepticism and have been difficult to reproduce (*18*, *19*). Thus, there exists a need to discover enzymes capable of C-C bond activation and cleavage required for deconstruction of non-hydrolysable plastics wastes into monomers or into bioavailable deconstruction products for upcycling (*9*).

Insect larvae, namely *Tenebrio molitor* (also known as the yellow mealworm), efficiently consume and degrade PE (*20*, *21*), PP (*22*), poly(vinyl chloride) (*21*), and polystyrene (PS) (*23*, *24*) at rates on the order of weeks — much faster than soil or marine environments (*25*) in which deconstruction typically takes tens to hundreds of years. Thus, the digestive systems of such insects and their gut microbiota may contain efficient enzymes for deconstruction of non-hydrolysable plastics; however, it remains unclear how much of this deconstruction is due to the activities of their gut microbiota or the host species itself (*20*, *26–29*). A recent report of PE deconstruction by the saliva of the greater wax worm suggested that deconstruction is initiated by the host (*16*), but the validity of these claims has been challenged (*18*). Most of the literature, though, hypothesizes that gut microbiota drive polymer deconstruction in insect systems through unvalidated oxidative mechanisms similar to alkane metabolism (*20*, *26*, *29–31*).

In this work, we sought to discover enzymes from yellow mealworm guts that participate in deconstruction of low-density PE (LDPE). LDPE was selected due to favorable material properties relative to other PEs, such as lower crystallinity and lower rigidity (*32*, *33*). Using a lower crystallinity substrate increases the chances of finding biological deconstruction agents, as amorphous domains are more accessible to enzymes and, thus, more readily bio-deconstructable (*34*). Previous studies have shown that yellow mealworms deconstruct LDPE, reducing weight-average molecular weight by approximately 61% and number-average molecular weight by approximately 40% (*20*). In this work, we identified yellow mealworm gut microbes and microbial enzymes that oxidize LDPE in isolation from their native environment by using Fourier transform infrared spectroscopy (FTIR) and x-ray photoelectron spectroscopy (XPS) analyses. More importantly, these studies revealed a sub-class of type I dye-decolorizing peroxidases (DyPs) capable of oxidizing LDPE films. Upregulation in gene count of DyPs in the gut microbiota of mealworms with plastic fed diets underscores the importance of these enzymes to initiating biodeconstruction. This enzyme sub-class contains a distinguishing hydrophobic loop insertion proximal to the active site that modulates the extent of oxidation and may participate in plastics binding. Surface residues proximal to this hydrophobic loop were found to be necessary for LDPE oxidation, suggesting a non-canonical route of substrate oxidation. Our study identifies a novel subclass of enzymes that play a pivotal role in initiating deconstruction of LDPE, opening new avenues for research into enzymatic polyolefin deconstruction and upcycling efforts.

## Results

### Yellow mealworm gut microbes oxidize low-density polyethylene

Yellow mealworms deconstruct LDPE, reducing its molecular weight by >33-62% upon ingestion (*20*, *35*, *36*). In the case of another nonhydrolyzable plastic, expanded polystyrene (EPS) foam, deconstruction appears to be driven by the yellow mealworm gut microbiota; plastic deconstruction is greatly reduced when mealworm guts are cleared of microbiota (*29*). Moreover, mealworms produce ^13^C labeled CO_2_ from ^13^C labeled PS only when microbes are present in their gut (*29*). As a direct result of these data and a growing number of reports of insect gut microbiota participating in LDPE deconstruction (*20*, *30*, *37*, *38*), we hypothesized that microbial enzymes in mealworm gut contents were responsible for LDPE deconstruction. We thus isolated microbes from the guts of mealworms fed diets of nonhydrolyzable plastics in the presence or absence of antimicrobial treatments (Fig. 1A). Using simple growth assays, we screened the LDPE deconstruction ability of over 300 isolates to reveal 21 taxonomically unique potential plastic degraders (Supplementary Table 1). Five top performing isolates, *Staphylococcus lentus, Enterococcus termitis, Corynebacterium variabile, Brevibacterium epidermidis,* and *Kocuria halotolerans* showed at least a 25% increase in optical density when grown in LDPE-containing mineral media compared to LDPE-free mineral media (Supplementary Fig. 1A). These strains were selected for further evaluation of their plastics deconstruction abilities.

**Figure 1:**
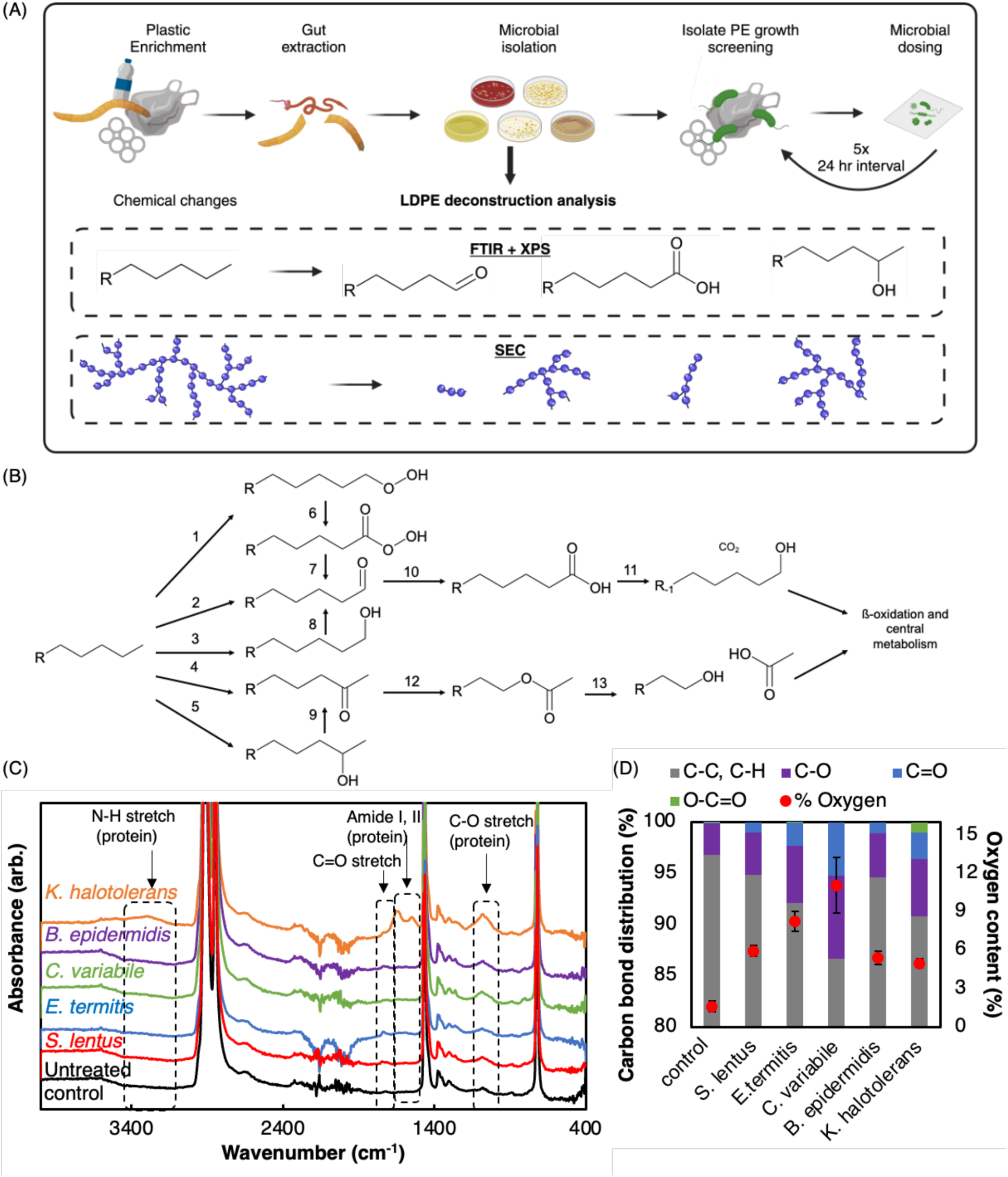
Yellow mealworm gut microbiota oxidize LDPE films. (A) Experimental procedure for microbial isolation, and materials characterization. Mealworms are fed LDPE to enrich their gut microbiome before microbial isolations from the gut were done aerobically, anaerobically, and in the presence of antibiotic and antifungal agents on several media. 335 total isolates were cultivated from PE enriched mealworm guts. (B) Hypothesized routes for enzymatic PE deconstruction. Each number represents a corresponding enzyme family: 1. Dioxygenase, 2. Peroxidase, 3. Monooxygenase, 4. Cytochrome p450 monooxygenase, 5. Monooxygenase, 6. Cytochrome p450 monooxygenase, 7. Esterase, 8. Alcohol dehydrogenase, Peroxidase 9. Alcohol dehydrogenase, Peroxidase, 10. Aldehyde dehydrogenase, 11. Decarboxylase, 12. Baeyer-Villiger Monooxygenase, 13. Esterase. (C) FTIR spectra of LDPE films after five 24-hour doses of mealworm gut isolates. (D) XPS spectral data detailing the C 1*s* bond distribution (main y-axis) and the extent of oxygenation of the PE films by the five isolates (secondary y-axis). Data represents the average of four technical replicates, taken at different locations on the same treated film. Error bars are the standard error of those four independent measurements.

The deconstruction of LDPE by mealworms and their gut microbiota is hypothesized to follow microbial alkane metabolism pathways, leveraging promiscuous enzymes that are active on hydrocarbons (*9*, *39–41*) (Fig. 1B). C-C and C-H bonds in LDPE films are proposed to be first enzymatically oxidized by monooxygenases, dioxygenases or peroxidases to generate primary or secondary alcohols. Further enzymatic oxidation of these primary and secondary alcohols produces aldehydes and ketones, respectively (*39*, *41*). In such a mechanism, the initial hydroxylation ‘activates’ the chain for deconstruction (the bond energy for a C-C bond is 607 kJ/mol compared to 314 kJ/mol in the CH_3_-CO bond (*42*)) and is subsequently hydroxylated in a second reaction, forming a geminal diol. The geminal diol then rapidly forms the observed ketone or aldehyde due to spontaneous, irreversible dehydration of the unstable diol intermediate (*43*, *44*). Further enzymatic oxidation and hydrolysis or decarboxylation are expected to produce fatty acids for beta-oxidation and/or carbon dioxide (Fig. 1B) (*9*).

We evaluated the propensity of top microbial isolates to deconstruct LDPE by evaluating the extent of oxidation on plastic substrates with diverse physicochemical properties (Table 1). Monocultures of the top five isolates were washed and then dosed daily on commercially available LDPE films over 5 days, adding fresh microbial inoculum to the film with each dose. The film was cleaned to remove bound proteins and characterized for chemical modification by X-Ray Photoelectron Spectroscopy (XPS) and Fourier Transform Infrared Spectroscopy (FTIR). All tested microbes increased the film oxygen content relative to untreated controls (one tail t-test, p<0.05), primarily in the form of carbonyl formation (C=O) (Fig. 1C–D). This carbonyl formation is consistent with proposed mechanisms of LDPE bio-oxidation (Fig. 1B, Supplementary Fig. 1B) and mimics chemical changes resulting from abiotic oxidation methods such as photo-oxidation (*45*) or cold plasma oxidation (*46*). *Corynebacterium variabile* treatment led to the largest increase in oxygen content relative to an untreated control (6.9-fold from 1.6% to 11%), followed by *Enterococcus termitis* (5.1-fold)*, Staphylococcus lentus* (3.7-fold)*, Brevibacterium epidermidis* (3.4-fold), and *Kocuria halotolerans* (3.1-fold) (Fig. 1D). FTIR spectra suggest that most isolates introduced aldehydes into PE chains by terminal oxidation (Figure 1C, Supplementary Table 2). However, microbes had distinct preferences for terminal or subterminal oxidation routes as inferred by the predominant detected C=O species (Figure 1B-D, Supplementary Table 2). In addition to new C=O bonds, treatment with each isolate resulted in minor increases in the C-O content of the film, likely as a result of the initial hydroxylation (Fig. 1D).

**Table 1:** Materials characteristics of plastic substrates used in this study.

| Plastic substrate | Form factor | $M_n$ (kg/mol) | $M_w$ (kg/mol) | Crystallinity |
| --- | --- | --- | --- | --- |
| Commercial Sigma Aldrich LDPE beads | Spherical bead (~1 cm) | 5.3 | 42.2 | 52.6% |
| Commercial Goodfellow LDPE | Film | 29.3 | 133.2 | 42.0% |
| In-house stripped LDPE | Film | 12.1 | 51.1 | 52.0% |

$M_n$ refers to the number average molecular weight
$M_w$ refers to the weight average molecular weight

Three isolates, *Corynebacterium variabile, Staphylococcus lentus*, and *Kocuria halotolerans* were chosen for SEC analysis due to different oxidation patterns on FTIR spectra (Supplementary Fig. 1C, Supplementary Table 2) that imply different enzymes responsible for oxidation events. Although isolated microbes induced chemical changes in the LDPE films, no change in the molecular weight distribution (MWD) of LDPE was detected via SEC (Supplementary Fig. 1C). However, due to the overlap between low molar mass products and solvent peaks in SEC chromatograms and the column set used, SEC is unable to detect and resolve soluble deconstruction products below ∼250 g/mol (Supplementary Fig. 2). Given the observed changes in LDPE film chemical modification (Fig. 1C) and LDPE-dependent growth of the microbial isolates (Supplementary Fig 1A), it is possible that some polymer chains are preferentially degraded into soluble products and metabolized upon initial oxidation, leaving high MW polymer chains for detection via SEC. An alternate interpretation, however, may be that efficient LDPE deconstruction is the result of synergistic action from a microbial consortium rather than isolates.

### Microbial communities in the mealworm gut initiate PE deconstruction via a secreted PE-oxidizing enzyme

We assembled and mined genomes for our top-performing microbial isolates to identify putative PE-oxidizing enzymes responsible for the observed changes in Figure 1C (Fig. 2A). All five isolate genomes were enriched relative to other species within their taxonomic Order in protein families proposed to oxidize alkanes such as dioxygenases, monooxygenases, and peroxidases (Fig. 1B, Supplementary Fig. 3, Fisher’s exact test; p-val < 0.05). These strains were likely present in the mealworm gut for their ability to metabolize lignin (*47–49*) and/or hydrocarbons such as cuticular waxes on leaves that form part of the mealworm diet (*50*). Thus, we hypothesized that these enzymes may have promiscuous activity on hydrocarbon-rich plastics.

**Figure 2:**
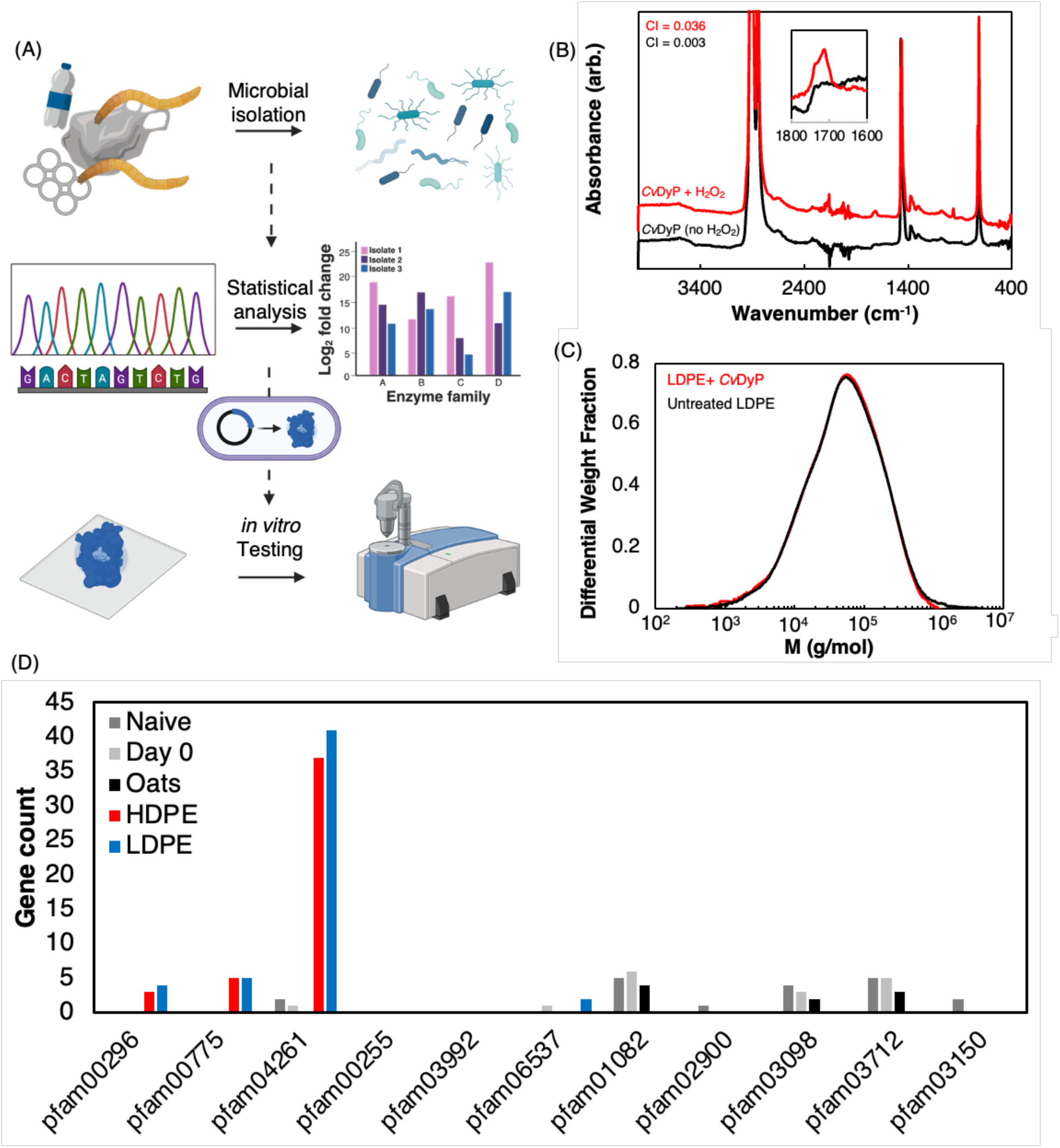
CvDyP oxidizes, but does not degrade, PE films. (A) Workflow for target enzyme identification in from mealworm gut isolates. Mealworm gut isolates were whole genome sequenced. Gene counts from each genome were compared with those of closely related taxa to determine families of enzymes from the classes outlined in Fig. 1B that may be more abundant in the mealworm gut isolates. Target enzymes were heterologously expressed into *E. coli* BL21 and tested *in vivo* on LDPE films. (B) FTIR data detailing oxidation of LDPE films by pure *Cv*DyP after washing with water and 70% ethanol. (C) SEC traces indicate that *Cv*DyP does not cleave C-C backbone bonds. (D) Prevalence of DyPs (pfam04261) in mealworm gut microbiomes fed LDPE and HDPE for 30 days relative to naïve (as they arrive in our laboratory), Day 0 (starved for 2 days), and oats (fed oats for 30 days) conditions. Protein families present in this analysis were selected as (1) putative enzymes that perform oxygenating chemistries (peroxidases, monooxygenases, dioxygenases) and that are (2) secreted.

The *Corynebacterium variabile* isolate encoded the highest diversity of putative PE-oxidizing enzyme families, which we evaluated for activity (Supplementary Fig. 3, Supplementary Table 3). Ten protein families (pfams), consisting of four classes of monooxygenases, three classes of dioxygenases, and three classes of peroxidases were identified as enriched in *C. variabile.* We randomly selected one representative enzyme from each of the ten pfams identified for *in vitro* testing on LDPE films (Supplementary Table 4). These enzymes were heterologously expressed in *E. coli* and lysates were used to treat LDPE films.

Among the tested enzymes, only the dye decolorizing peroxidase (DyP; pfam 04261), denoted *Cv*DyP (IMG gene ID Ga0530663_0293_23766_25013), chemically modified LDPE films and acted as a LDPE oxidase (Fig 2B). Three 90-minute treatments of LDPE films with purified *Cv*Dyp and cofactor H_2_O_2_ led to the formation of ketones and aldehydes, evidenced by FTIR spectral peaks at 1710 cm^-1^ and 1740 cm^-1^, respectively. Carbonyl peaks were confirmed to be a direct result of enzymatic activity, as the peak persists after washing with water and ethanol. Washing is critical for accurate identification of oxidation as it removes surface-bound protein (*18*) that generates a confounding signal in the carbonyl region(Supplementary Fig. 4). To quantify this activity, we calculated the carbonyl index (CI) or ratio of the maximum peak height between 1700– 1745 cm^-1^ and the maximum peak height between 1400–1500 cm^-1^ (*51*) (Supplementary Fig. 5). Enzyme treatment increased film CI by more than 10-fold relative to a H_2_O_2_-free, inactive *Cv*DyP control (Fig. 2B). Sequence analysis of the enzyme with SignalP confirmed that this DyP was secreted by its microbial host and peroxidase activity was confirmed using model substrate pyrogallol (Supplementary Fig. 6). Despite the ability of *Cv*DyP to oxidize PE films, it is unable to cleave C-C bonds in PE after 20x 90-minute treatments (Fig. 2C).

DyPs are absent in the yellow mealworm genome but are highly abundant in the metagenomes of mealworm gut microbial communities fed on LDPE (41 gene counts) and high-density PE (HDPE) (37 counts) relative to oats fed (0 counts) mealworms (Fig. 2D). Since DyPs are present only in the gut microbiome and not the host, our work suggests that gut microbial communities may play an important role in LDPE deconstruction in yellow mealworms (*52*). DyP producing microbes are far more prevalent in plastic-containing guts, implying that this oxidative event is critical for the LDPE deconstruction process.

### PE-oxidizing DyPs are ubiquitous

As DyPs are ubiquitous in bacteria and fungi, we investigated whether there were unique sequence and structural features that dictate if a DyP can function as an LDPE-oxidase. DyPs are divided into three major classes based on size: i) P or *primitive* DyPs (<300 amino acids); ii) I or *intermediate* DyPs (300–400 amino acids); and iii) V or *advanced* DyPs (>400 amino acids) (*53*, *54*). *Cv*DyP is a class I (subclass I3) DyP. Thus, we heterologously expressed and tested 11 enzymes across all four sub-classes of class I DyPs, both native and external to the yellow mealworm gut microbiome to determine if LDPE oxidation is a general property of class I DyPs, or the I3 subclass of DyPs (Supplementary table 5). DyPs across all tested sub-classes oxidize LDPE as measured by an increase in CI; only *Bl*DyP and *Bo*DyP did not show activity over the *E.coli*-noDyP control (Supplementary Fig. 7). Thus, PE-oxidizing DyPs are not restricted to the guts of yellow mealworms and may be found in a number of environments including well water (*55*), cheese rind metagenomes (*56*), and stool samples (*57*, *58*).

To ensure we identified enzymes that act on PE chains rather than potential additives, we tested purified class I3 DyP (*Cv*DyP, *Cn*DyP, and *Cs*DyP) activity on LDPE films that were stripped of additives in-house *via* Soxhlet extraction (Fig. 3a, Table 1, Supplementary Fig. 8); a list of stripped additives identified *via* GC-MS can be found in Supplementary Data File 2. *Cn*DyP and *Cs*DyP were selected for further study as the nearest phylogenetic neighbors of *Cv*DyP. Again, each of these enzymes was confirmed as a peroxidase by demonstrating activity on model substrate pyrogallol (Supplementary Fig. 9a). All three enzymes were active on stripped LDPE, meaning that oxidation occurs on the polymer chain, not on any additives (Fig. 3a). However, quantitative performance is substrate specific. Relative enzyme activity on stripped LDPE (∼83 kg/mol, Fig. 3A) differs from commercial LDPE (∼133 kg/mol, Supplementary Fig. 9b)., underscoring the importance of polymer characteristics for enzyme performance Nonetheless, our results indicate that the I3 class of DyPs both native and not native to the mealworm gut are able to oxidize PE chains in LDPE films.

**Figure 3:**
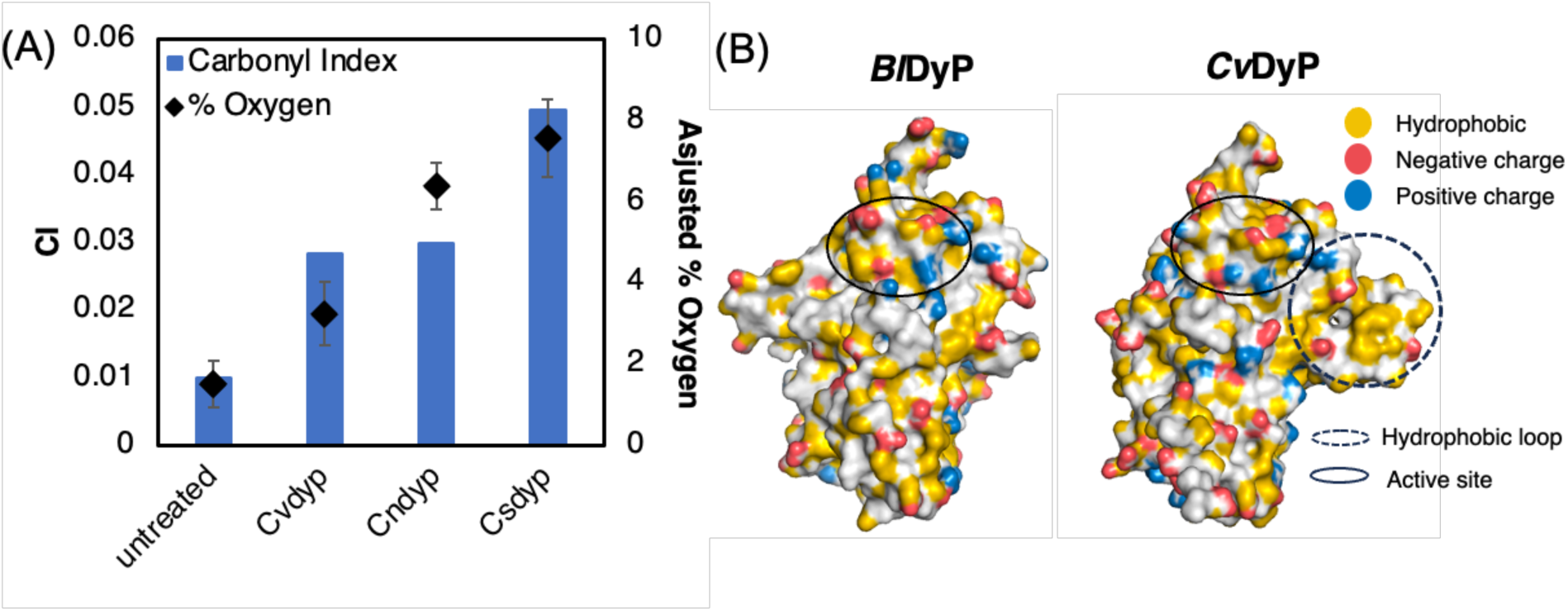
Purified type I3 DyPs oxidize PE films. (A) CI from FTIR spectra (calculated as the ratio of the maximum peak height between 1700–1745 cm^-1^ and of the maximum peak height between 1400–1500 cm^-1^) and % oxygen on stripped LDPE films after treatment with purified DyP peroxidase samples and after washing with water and 70% ethanol. (B) YRB hydrophobicity maps of select inactive (*Bl*DyP) and active (*Cv*DyP) detailing hydrophobic atoms in yellow, positively charged atoms in blue, and negatively charged atoms in red.

### A prominent hydrophobic loop near the active site modulates DyP PE-oxidase activity

We hypothesized that conserved structures across LDPE-active DyPs may be important for LDPE oxidation. Thus, we predicted the structure of each tested DyP via AlphaFold 2.0 (all with pLDDT confidence scores >88.8) and used them in a multiple structural alignment (*59*). Secretion signal peptides were identified via SignalP version 6.0 and removed prior to structural simulations to aid alignment. LDPE-active DyPs contain a unique loop region extending outward from the body of the protein and proximal to the predicted active site (Fig. 3B, Supplementary Fig. 10) that deviates from non-PE oxidizing DyPs. Importantly, the loop is hydrophobic, ranging from approximately 33% (*Bl2*DyP) to 54% (*Cn*DyP) of residues with hydrophobic side chains. This loop in *Cv*DyP is encoded by: YNYDLPVTPSSADALVDADPVALSDT, with YNY residues at the start of the sequence and GL residues at the end of the sequence being conserved across all tested type I DyPs (Supplementary Fig. 11). LDPE oxidase activity directly correlates with hydrophobic loop length and hydrophobicity (Supplementary Fig. 7, 11); the enzymes with the longest and most hydrophobic loops are the top performing LDPE oxidases. Non-PE active enzymes contain a much smaller 12 amino acid loop that is more hydrophilic (*60*) (Fig. 3B, Supplementary Fig. 10-12). The extension of the hydrophobic loop outward from the center of the protein and its proximity to the predicted active site (*61*) suggests that the hydrophobic loop may be important for productive binding of highly hydrophobic PE chains in orientations favorable for catalysis.

Mutant DyPs were generated to validate the role of the hydrophobic loop for LDPE-oxidase activity. The four tested DyPs from *Corynebacterium* strains (*Cp*DyP, *Cv*DyP, *Cs*DyP, and *Cn*DyP) were selected as engineering targets due to their high sequence homology but differing hydrophobic loop structures and lengths. The smaller, less hydrophobic loop region from *Cp*DyP was swapped with the hydrophobic loops in *Cv*DyP, *Cn*DyP, and *Cs*DyP to create mutant hydrophobic loop reduction and extension chimeras. Reducing the hydrophobic loop size and hydrophobicity reduced LDPE-oxidase activity relative to the wild type (Fig. 4). Similarly, inserting the longer, more hydrophobic loops from *Cv*DyP, *Cn*DyP, and *Cs*DyP into *Cp*DyP enhanced LDPE-oxidase activity over the wild type *Cp*DyP (Fig. 4). Quantifying the hydrophobicity of these mutants around the active site and loops via a sequence hydrophobicity index revealed a positive correlation with observed LDPE activity (R^2^ = 0.43; Supplementary Fig. 13). These results confirm that this divergent hydrophobic loop region has a significant role in DyP LDPE activity.

**Figure 4:**
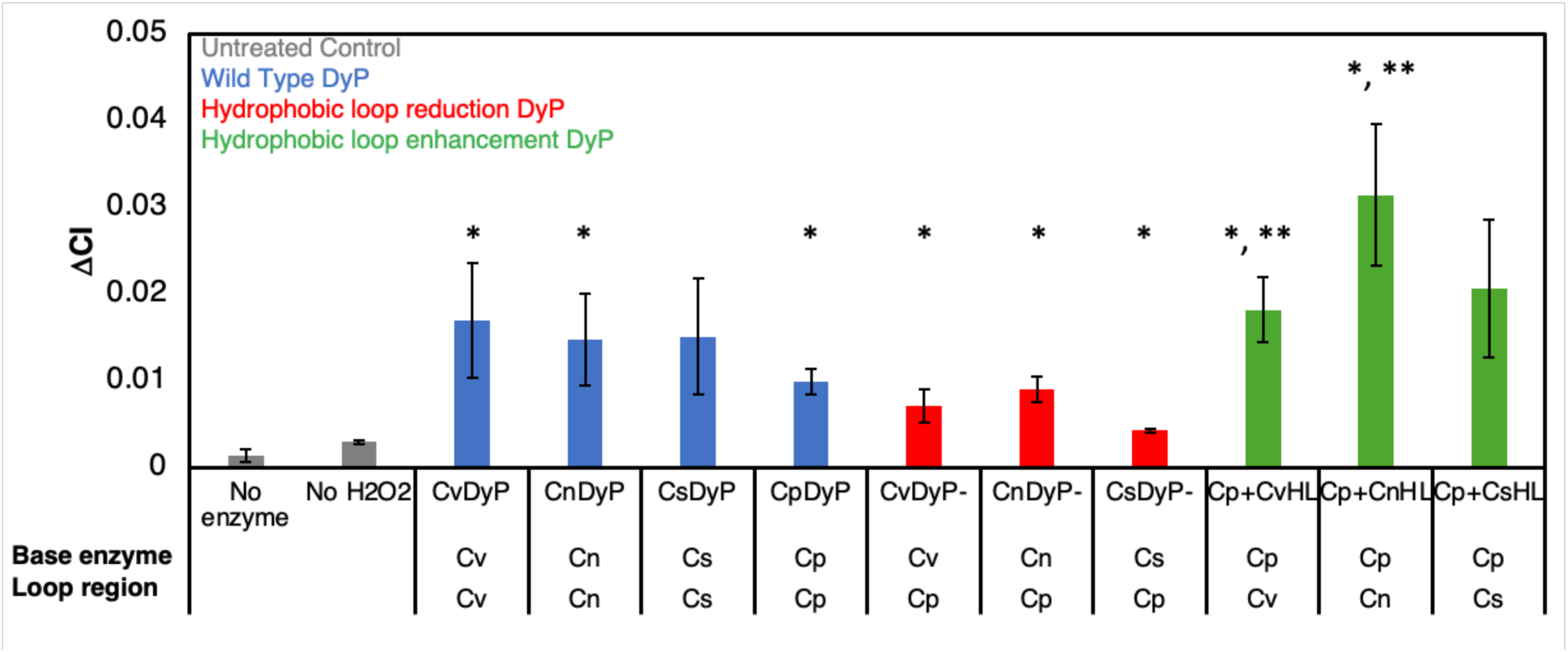
The hydrophobic loop region in DyPs can be used to modulate PE-oxidase activity. Changes in carbonyl index (Λ1CI) observed between stripped LDPE films before and after enzymatic treatment with purified enzyme, detailing LDPE-oxidase activity by loop-engineered DyPs relative to native DyPs. Films were washed with water and 70% ethanol prior to FTIR spectrum collection. xxDyP-nomenclature refers to the base (xx) enzyme hydrophobic loop being removed for the smaller, less hydrophobic loop of CpDyP. Cp+xxHL nomenclature refers to the addition of the hydrophobic loop from xxDyP to CpDyP in place of the native hydrophobic loop. Error bars represent standard error across four independent biological replicates. *Statistically significant increase in ΔCI relative to no enzyme control via one-tailed t-test, p<0.05. ** Statistically significant increase in ΔCI relative to *Cp*DyP via one-tailed t-test, p<0.05.

### Surface-exposed tryptophans catalyze non-canonical oxidation of LDPE chains external to the protein

We next tried to determine how DyPs non-terminally oxide PE substrates as they are unable to fit within the canonical active site. However, studies with reactive blue (RB19), a bulky dye that cannot fit into the DyP active site, demonstrate a non-canonical oxidation mechanism mediated by surface-exposed aromatic residues that can harbor radicals (*62*, *63*). These radicals allow for oxidation of the otherwise sterically hindered RB19 by facilitating long chain electron transport from the substrate to the heme cofactor (*62*, *63*) and are conserved across type I DyPs;using *Tc*DyP as a basis, W263 is conserved and W356 is conserved as a tryptophan or a tyrosine subclass-wide (*63*)(*63*). Therefore, we hypothesized that LDPE oxidation occurs *via* the same mechanism and identified W231 and W356 in *Cv*DyP as analogous residues to W263 and W376 that are necessary for RB19 oxidation by *Tc*DyP (*63*) (Fig. 5a).

**Figure 5:**
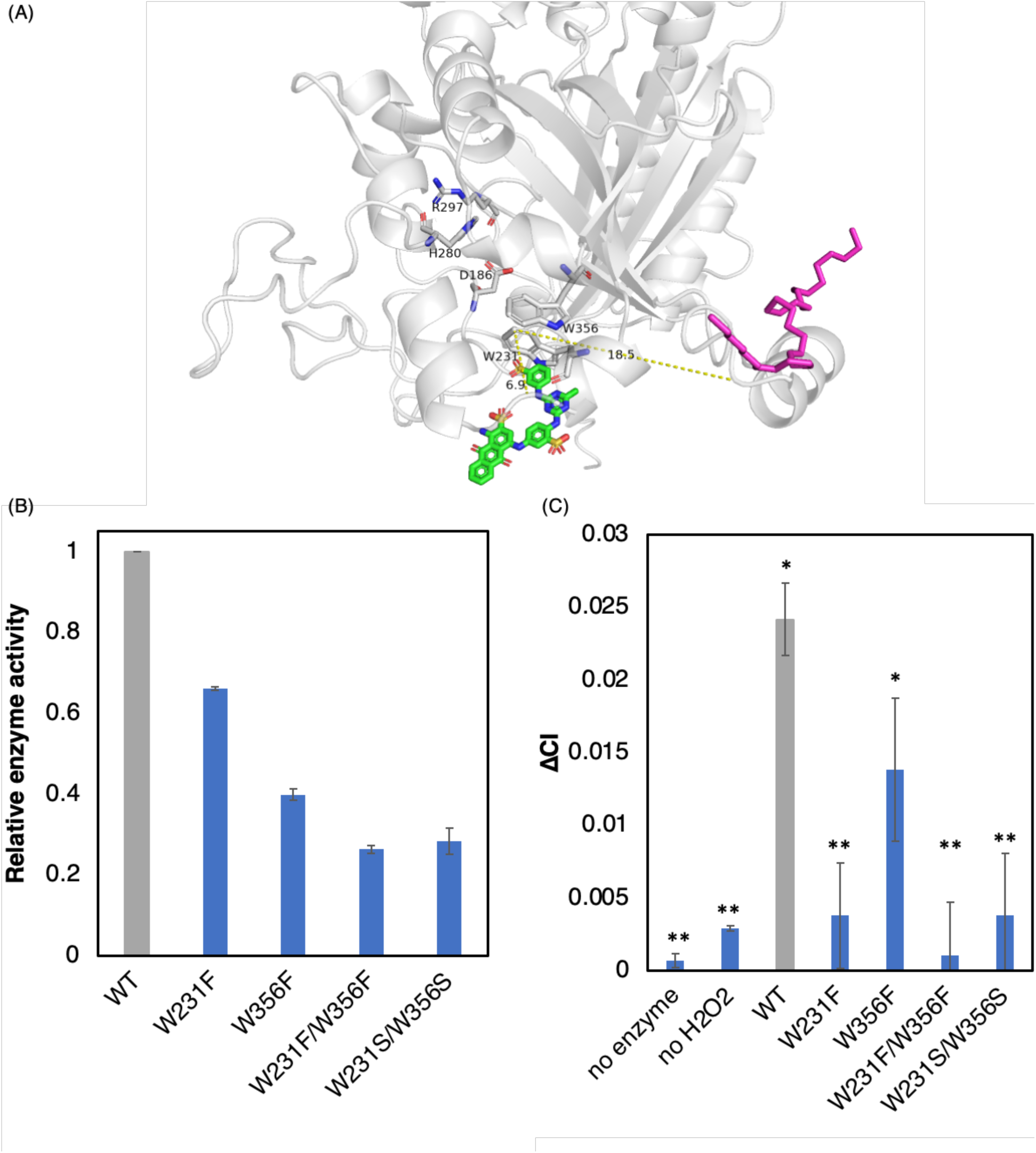
Surface exposed aromatic residues permit LDPE oxidation. (A) Pymol rendering of Alphafold predicted CvDyP structure with active site (D186, H280, and R297) and surface exposed aromatics (W231 and W356) displayed as sticks. Distances between LDPE-oxidizing W231 and bulky aromatic substrate reactive blue 2 (green) and model alkane dotriacontane (pink) are displayed. (B) Relative enzyme activities of CvDyP mutants on model substrate pyrogallol. (C) Change in carbonyl index of CvDyP and mutants after treatment on stripped LDPE. *Statistically significant increase in ΔCI relative to no enzyme control via one-tailed t-test, p<0.05. **Statistically significant difference in ΔCI relative to WT via one-tailed t-test, p<0.05.

To test this hypothesis, we mutated these two tryptophans conservatively to non-radical forming phenylalanine or serine known to preserve the environment around the heme cofactor (*62*, *63*). Control studies with model substrate pyrogallol, which binds the canonical active site, demonstrate that all mutants (W231F, W356F, W231F/W356F, and W231S/W356S) retained canonical peroxidase activity (Fig 5b). However, there was a 40-70% decrease in activity relative to wild type, perhaps due to minor conformational shifts in the protein structure and/or loss of non-canonical surface activity. All W231 mutants completely lost LDPE oxidase activity relative to wild type control, demonstrating that it is necessary for DyP activity on plastic substrates (Figure 5c). W356 is involved in plastics deconstruction although it is not essential for activity; W356 mutants lost 48.5% of activity relative to wildtype on PE substrates (Figure 5c). These results are consistent with our hypothesis and suggest that LDPE oxidation is catalyzed by surface-exposed amino acid (W231 and W3566) residues on DyPs.

Our observations regarding the surface catalysis of PE substrates (Fig. 5) and the importance of a hydrophobic loop for activity (Fig. 4) suggest a model for LDPE catalysis *via* DyPs. LDPE oxidation proceeds *via* the surface-exposed radical mechanism only, as it cannot bind the relatively small canonical active site, and the hydrophobic loop aids with substrate binding and positioning against these residues for activity. Removing this loop reduces PE interaction with these surface residues and thus reduces activity (Fig 4). However, this loop-mediated mechanism is only possible for substrates of a minimum size, as the substrate must be large enough to bind to the hydrophobic loop and reach the surface oxidation site. Model alkanes (C_10_–C_40_) that failed to dock to the canonical active site, as simulated by Autodock, were not oxidized by top performing DyP mutant, Cp+CNHL DyP (Supplementary Fig. 14). Moreover, C_32_ alkane dotriacontane, the largest substrate we could simulate with off-the-shelf tools, is too far from W300 and Y424, analogous to *Cv*DyP W231 and W356, to be oxidized in the same manner bulky dyes are oxidized when its docking was forced to the *Cp+CNHL*DyP hydrophobic loop using Autodock (Supplementary Fig. 15). That is, surface-mediated activity is only possible when the substrate is large enough for the hydrophobic loop to correctly position it, such as PE chains and their deconstruction products. Under optimum conditions, dotriacontane (C_32_) binding was calculated to be ∼9.5 Å short from interacting with the surface-exposed radicals for catalysis (Supplementary Fig. 15). By approximating a C-C bond length of 5 Å per six carbons estimated by Pymol, the C_32_ alkane was found to be short ∼12 carbons. In other words, a bulky substrate must be at least 44 carbons long to be correctly positioned by the hydrophobic loop to access the surface residues. This nonconservative estimate of the minimum substrate size is suggested by Autodock simulations as the specific position at which a PE chain binds to the hydrophobic loop. The orientation and branching density of bound PE chains and the flexibility of the hydrophobic loop region all impact actual binding.

## Discussion

In this work, we identified type I DyPs as a class of LDPE-oxidizing enzymes and provide evidence for their role in LDPE deconstruction in yellow mealworms by showing an enrichment in gene count of DyPs in PE-fed mealworms relative to non-plastic-fed controls. DyPs initiate LDPE deconstruction by oxidizing PE chains to form aldehydes or ketones. Rigorous controls and characterization confirmed the validity of these observations and reduced false positives in activity measurement. LDPE cleaning prior to materials characterization allowed for distinction between biological contaminants and true chemical modification. LDPE substrates stripped of additives were similarly oxidized, demonstrating enzymatic activity on the polymer backbone, rather than polymer additives. Importantly, evidence of LDPE oxidation is absent on H_2_O_2_-free controls, confirming peroxidase activity. The reliability of these findings was demonstrated by independent quantitative spectroscopic techniques, FTIR and XPS. We identified a divergent loop characteristic of LDPE-active DyPs and demonstrated that its hydrophobicity can be tuned to enhance LDPE activity. Lastly, we provide support for a non-canonical mechanism of LDPE oxidation through solvent-exposed, radical-harboring amino acid residues by abolishing LDPE oxidation upon mutation of putative oxidative residue W231.

This work suggests that fully biological routes for LDPE deconstruction are feasible by confirming enzymatic oxidation of PE chains. Abiotic pre-treatments such as chemical treatment (*64*) or thermal oxo-deconstruction were previously thought to be necessary to make plastics chemically available for biodeconstruction (*65*). Though such an oxidative step has been hypothesized as the first step in enzymatic LDPE deconstruction (*9*), this step had not been directly demonstrated until now. Activity of LDPE deconstructing enzymes such as laccases, manganese peroxidases, and alkane hydroxylases are limited to pre-oxidized or very low molecular weight (M_w_ ∼4 kg/mol, M_n_ ∼1.7 kg/mol) polymers (*17*, *66–68*). Our data conclusively show that abiotic oxidation or pretreatment is not required for biological deconstruction and that biological activation chemistries do exist.

Phenol oxidases (*16*) and multi-copper oxidases (*69*) have been reported to oxidize and deconstruct non-pretreated LDPE. Both reports show carbonyl formation after enzyme treatment *via* FTIR or Raman spectroscopy, consistent with the oxidation performed by DyPs, but neither study leveraged inactive enzyme controls (*16*). This lack of controls, coupled with a lack of repeatability has led the phenol-oxidase study to be scrutinized (*18*), as the possibility of non-specific enzyme binding to the plastic leading to the formation of new carbonyl peaks cannot be ruled out. We confirm that new carbonyl groups are a result of DyP activity rather than non-specifically bound protein by including a DyP control in the absence of H_2_O_2_ and by including FTIR spectra before and after washing LDPE films. Additionally, any oxidative or deconstructive activity by phenol oxidases or multi-copper oxidases on LDPE could be the result of activity on additives found in the plastic. We confirmed here that DyP activity occurs directly on LDPE chains by stripping the plastic of additives prior to its use. Both of the aforementioned studies also report medium chain ketones and/or carboxylic acids as deconstruction products *via* gas chromatography, but these compounds may have been inherent to the plastic and leached out into the reaction mixture, a possibility that cannot be ruled out due, again, to the lack of inactive enzyme controls in this study. Moreover, the generation of medium chain deconstruction products implies that the single purified enzyme performs oxidation and cleavage of C-C bonds, which is highly unlikely for a single enzyme, as the chemistry for C-C oxidation is far different than that needed to cleave C-C bonds in the PE backbone (*9*, *18*). By confirming that oxidation occurs directly on PE chains, we definitively show that DyP peroxidases act as a first enzymatic step in that series by oxidizing the PE chains, priming them for deconstruction.

DyPs are known for their substrate promiscuity and propensity to oxidize bulky polymeric substrates such as lignin (*53*, *70–72*) and have been shown to oxidize and degrade UV-oxidized PS (*73*). Oxidation of substrates that cannot fit in the active site is made possible by catalytic radicals on surface exposed, aromatic residues (*62*, *74*), but the importance of these residues has only been demonstrated on the oxidation of bulky dyes such as RB19 (*62*, *63*). We confirmed the importance of W231 in *Cv*DyP, showing that it must be present to oxidize LDPE. This finding is consistent with the mechanism for RB19, as W231 is analogous to W263 of *Tc*DyP, the residue responsible for RB19 oxidation (*63*). Therefore, we propose that surface residue W231 serves as a non-canonical oxidation site for LDPE chains. This non-canonical oxidation is enhanced by a hydrophobic loop region proximal to the solvent-exposed aromatics, where the loop region can act as an anchor and bind to the extremely hydrophobic LDPE, making LDPE chains sterically available for protein radicals that facilitate the transfer of electrons and LDPE oxidation. Thus, the hydrophobic loop and surface aromatic regions are promising targets for engineering to enhance LDPE oxidation.

DyPs provide a crucial first step in the LDPE biodeconstruction process through oxidation of PE chains. Due to the diverse nature of this enzyme class (*53*, *54*), there may be DyPs from sources other than mealworm guts that are more effective at PE oxidation than those reported in this study. Moreover, studying the oxidation mechanism of DyPs is essential to improve LDPE-oxidase activity. Such activities are informed by the identification of the hydrophobic loop region and residue W231 in this work. These findings can be further developed to create design rules for LDPE-oxidizing DyPs and to engineer LDPE-oxidases with improved activity or tolerance to industrial processing conditions. By providing the first step in biological LDPE deconstruction, this study provides a route to discover enzymes downstream of DyPs in the deconstruction pathway. Ultimately, this knowledge can enable the development of sustainable, biologically driven approaches for plastics deconstruction.

## Materials and Methods

### Mealworm cultivation

For each gut cultivation experiment, 100 yellow mealworms (Medium yellow mealworms were purchased from Rainbow Mealworms. Compton, CA, USA) were starved for 48 hours upon arrival and then placed in a Tupperware box with approximately 0.5 g of LDPE in a static incubator at 25 °C, 70% humidity. Control worms fed oats were given 1.0 g of oats every 3 days to supplement feeding. In order to avoid beetles and mealworm cannibalism, dead and pupated larvae were removed every 2 days.

### Fourier transform infrared spectroscopy FTIR

Commercial (0.035 mm thick, biaxially oriented LDPE films, purchased from Goodfellow Cambridge Limited, Huntingdon, England. Catalog number LS580568) LDPE films and in-house stripped LDPE films were analyzed for changes to the inherent functional groups using FTIR (Thermo Scientific Nicolet iS5 FTIR Spectrometer, Pittsburgh, PA). Spectra for each powdered sample were recorded on a diamond crystal, attenuated total reflectance (ATR) cell. Spectra were recorded in the range of wavelengths 4000– 500 cm^−1^ with a minimum of 32 scans and a spectral resolution of 0.482 cm^−1^. Films were rinsed with 500 µL of water followed by 500 µL of 70% ethanol by vortexing for 1 minute at 200 rpm. If FTIR spectra indicated microbial or protein contamination, 70% ethanol on a cotton swab was lightly wiped over the surface to clean residual contamination.

### Size Exclusion Chromatography (SEC)

Molecular weight characterization through SEC was conducted on a TOSOH HLC-8312GPC/H with two TSK_gel_ GMHHR-H(20)HT columns and one TSK_gel_ G2000H_HR_ in series, coupled with refractive index (RI) and viscosity detectors. HPLC-grade 1,2,4-trichlorobenzene (TCB) stabilized with 500 ppm butylated hydroxytoluene (BHT) was used as the mobile phase. LDPE samples (3–10 mg) were dissolved in mobile phase (3–6 mL) at 140 °C for a minimum of two hours to generate nominal concentrations of 2 mg/mL. 300 µL of samples were injected and eluted for 80 minutes at a flow rate of 0.8 mL·min^-1^ at 140 °C.

MWDs were determined using the RI responses, and SEC samples were calibrated against linear polystyrene standards (12 runs between 6×10^2^–2×10^6^ g/mol^-1^; Supplementary Figure 2). Molar masses of LDPE obtained were corrected using the Mark-Houwink relationship (following previously detailed methods) (*75*).

### Metagenomic DNA extraction and sequencing

Following mealworm cultivation, guts from LDPE-fed, HDPE-fed, and oats fed guts, along with naïve (as they arrive from Rainbow Mealworms) and starved (starved for 2 days after arrival from Rainbow Mealworms) were extracted from 10 mealworms, pooled together, and vortexed for 5 minutes at 2000 rpm to extract gut contents. A quick ∼5 second centrifuge step was then used to pull remaining gut tissue to the bottom of a microcentrifuge tube without disturbing microbial biomass. The supernatant containing microbial content was removed and DNA was extracted using Monarch Genomic DNA extraction kit and sent to the Joint Genome Institute (JGI) for sequencing. An input of 50 ng of genomic DNA was sheared to 6 kb – 10 kb using the Megaruptor 3 (Diagenode). The sheared DNA was treated with exonuclease to remove single-stranded ends, DNA damage repair enzyme mix, end-repair/A-tailing mix and ligated with amplification adapters using SMRTbell Express Template Prep Kit 2.0 (PacBio) and purified with ProNex Size-Selective Purification System (Promega). The purified ligation product was split into two reactions and enriched using 10–18 cycles of PCR using SMRTbell gDNA Sample Amplification Kit (PacBio). The amplified product was combined and treated with DNA damage repair enzyme mix, end-repair/A-tailing mix and ligated with barcoded overhang adapters. Up to sixteen libraries were pooled in equimolar concentrations and the pooled libraries were size-selected using the 0.75% agarose gel cassettes with Marker S1 and High Pass protocol on the BluePippin (Sage Science). PacBio Sequencing primer was then annealed to the SMRTbell template library and sequencing polymerase was bound to them using Sequel II Binding kit 2.0. The prepared SMRTbell template libraries were then sequenced on a Pacific Biosystems’ Sequel IIe sequencer using SMRT Link 10.2, tbd-sample dependent sequencing primer, 8M v1 SMRT cells, and Version 2.0 sequencing chemistry with 1×1800 sequencing movie run times.

Metagenomes sequenced by the JGI are publicly available on Integrated Microbial Genomes & Microbiomes (IMG/MER) under the following IMG Genome portal numbers:

Naïve mealworm gut metagenome: 3300066518

Day 0 (starved) mealworm gut metagenome: 3300066519

30-day HDPE fed mealworm gut metagenome: 3300056814

30-day LDPE fed mealworm gut metagenome: 3300056790

30-day oats fed mealworm gut metagenome: 3300066518

### Metagenomic data analysis to identify protein families of interest

All genes corresponding to pfams containing function peroxidase, monooxygenase, dioxygenase, and laccase were identified using the IMG Genome Function Profile tool across the metagenomes. These Genes obtained from JGI sequenced metagenomes were screened then with signalP 6.0 Fast version to determine secretion tag presence. Signal P settings were organism-other, and format none. All genes that SignalP tagged as having any output besides “other” were considered to be secreted. Any gene without a secretion tag was eliminated from a .gff annotation file for pfams generated by the JGI standard pipeline, and then the gene count of each pfam was used for plotting.

### Microbial isolation from mealworm guts

Mealworms were fed diets of HDPE, LDPE, PP, and PS with or without oats, according to the mealworm cultivation method listed above. Mealworms were enriched on plastic for 20 days, with gut extractions and isolations occurring on days 5, 10, 15 and 20. Mealworm guts were extracted according to protocol in mealworm cultivation section. One mL of sterile phosphate buffered saline (PBS) was added to each sample and samples were vortexed for 5 minutes at 2000 rpm. Mealworms gut solutions were plated onto the following media agar: LB (Sigma-Aldrich Catalog No. L3147-1KG), YPD (Sigma-Aldrich, Catalog No. Y1500-250G), PDA (Millipore Sigma, Catalog No. 110130), Sheep’s Blood (Rockland Chemical Company, Item No. R111-0050) (*76*), and MacConkey agar plates (Becton, Dickinson and Company, SKU: 211387). Mealworm guts were also plated onto each type of agar including 50 mg/mL of antibiotics penicillin and streptomycin or including 50 mg/mL of antimycotic amphotericin. Individual colonies from morphologically unique isolates were picked, grown in the corresponding liquid medium, and stocked into –80 °C storage in 20% glycerol. Starting with mealworm gut extractions, this process was repeated anaerobically to cultivate any anaerobic microbes from the mealworm gut.

### Growth assay screening for PE deconstruction

All cultivated microorganisms from mealworm gut isolations were screened for PE deconstruction ability through a growth assay. Microbes were grown in a mineral medium (MM) (1 g NaH_2_PO_4_, 0.25 g MgSO_4_ * 7H_2_O, 0.1 g KH_2_PO_4_, and 0.5 g of yeast extract per 500 mL water) as a control, and then with 0.3% w./v.LDPE powder (LDPE powder was purchased from Goodfellow Cambridge Limited, Huntingdon, England; catalog number LS563303). Microbes that showed improved growth after 7 days on MM + LDPE relative to MM via OD 600 measurement were selected for further study. The mineral medium was bootstrapped with a low concentration of yeast extract to allow microorganisms to adapt and produce the enzymes needed to grow on PE as a primary carbon source. Microbial isolates that showed an increase in growth on MM + LDPE relative to MM were selected as potential plastic degraders.

### Plastic oxidation assays by microbial isolates

Starter cultures of each isolate were grown in Tryptic Soy Broth (Sigma-Aldrich, Catalog No. 22092-500G) from freezer stocks. Optical densities were recorded of each overnight culture and the number of microbes for inoculation was normalized accordingly, assuming OD600 as a proxy for number of bacterial cells. 3 mL of OD 1.0 overnight culture was pelleted, washed twice using MM and then resuspended in MM for inoculation. MM was used to mimic conditions used in growth screening assays and to provide microorganisms with non-carbon-based essential nutrients. Inoculum was dosed onto an MM plate, covered with an LDPE film (Goodfellow Cambridge Limited, Huntingdon, England. Catalog number LS580568), and then a second inoculum was dosed atop the film. Following the same protocol, microbes were dosed onto films daily for 5 days. After treatment, films were washed via vortexing in 500 µL water for 2 minutes at 2000 rpm, followed by vortexing in 500 µL of 70% ethanol under the same conditions. Films were then subject to FTIR and X-Ray Photoelectron Spectroscopy (XPS) analysis. Films subject to XPS analysis were additionally washed in Sodium Dodecyl Phosphate (SDS) to remove additional adventitious carbon contaminants. Additionally, films were analyzed via SEC to determine changes in the MWD of the polymer to delineate which isolates can degrade LDPE.

### XPS

XPS spectral scans, C1s, and O1s scans were collected on four locations on each film, using the following settings: C1s spectra were analyzed assuming a C-C, C-H peak at 285.0 eV and then peak shifts of 1.5 eV, 3.0 eV, and 4.5 eV for C-O, C=O, and O-C=O peaks, respectively (*77*). All peak fitting analysis was performed using Casa XPS. In survey spectra that contained residual contamination in the form of nitrogen or sodium from protein or SDS from washing, residual oxygen was subtracted by subtracting 1% O for every 1% N and by subtracting 1% O for every 4% Na, respectively. All spectral analysis was assessed by averaging the results from survey scans on four randomly selected points each film.

### Microbial isolate DNA extraction and genome sequencing

Microbial isolates *Staphylococcus lentus, Enterococcus termitis, Corynebacterium Variabile, Brevibacterium Epidermidis,* and *Kocuria halotolerans* were grown overnight in TSB and DNA was extracted using NEB Monarch Genomic DNA Purification Kit (Monarch Genomic DNA purification Kit #T3010L, New England BioLabs, Ipswich, MA, USA.), following the kit protocol. Genomic DNA quality was confirmed via qubit and DNA gel electrophoresis. Genomic DNA was sequenced by the Joint Genome Institute.

### Isolate Minimal Draft Genome Sequencing and Assembly

The draft genome of each isolate *Staphylococcus lentus, Enterococcus termitis, Corynebacterium Variabile, Brevibacterium Epidermidis,* and *Kocuria halotolerans* was generated at the DOE Joint Genome Institute (JGI) using Illumina technology (*78*). An Illumina standard shotgun library was constructed and sequenced using the Illumina NovaSeq S4 platform which generated at least 6.7 million reads for each isolate, and completeness of 98.42 – 100%. Raw Illumina sequence was quality filtered using BBTools (*79*) per SOP 1061. The following steps were then performed for assembly: (1) artifact filtered and normalized Illumina reads were assembled with SPAdes (version v3.15.3; – phred-offset 33 –cov-cutoff auto -t 16 -m 64 –careful -k 25,55,95) (*80*); (2) contigs were discarded if the length was <1kb (BBTools reformat.sh: minlength=1000 ow=t).

Genomes sequenced by the JGI are publicly available on Integrated Microbial Genomes & Microbiomes (IMG/MER) under the following IMG Genome portal numbers:

*Brevibacterium epidermidis*: 8012935351

*Corynebacterium variabile*: 8081958687

*Enterococcus termitis*: 8012939035

*Staphylococcus lentus*: 8012942269

### Genome annotation

All genomes were annotated using annotation System IMG (IMGAP v5.1.13). Gene calling program was GeneMark.hmm-2 v1.25_lic; INFERNAL 1.1.3 (Nov 2019); Prodigal v2.6.3; tRNAscan-SE v.2.0.12 (Nov 2022). The annotation algorithm was: lastal 1256; HMMER 3.1b2; signalp 4.1; decodeanhmm 1.1g. The following databases were used to annotate each gene: Rfam 13.0; IMG-NR 20211118; SMART 01_06_2016; COG 2003; TIGRFAM v15.0; SuperFamily v1.75; Pfam v34.0; Cath-Funfam v4.

### Comparative genomic analysis for enzyme selection

Genomic sequences of each isolate were mined for protein families (pfam) of monooxygenases, dioxygenases, laccases, and peroxidases using the JGI Integrated Microbial Genomes (IMG) tool (*81*). The raw pfam gene counts in the microbial genomes were compared to the average gene count of the same pfam across the taxonomic order. The Joint Genome Institute (JGI) integrated microbial genomes (IMG) (*81*) workspace was used to compile all genomes from the taxonomic order of each strain *Staphylococcus lentus, Enterococcus termitis, Corynebacterium variabile, Brevibacterium epidermidis,* and *Kocuria halotolerans*. The IMG Genome Function Profile tool was then used to generate a count of all genes corresponding to putative plastic degrading pfams (Supplementary Table 3) across the genome in the taxonomic order. These jobs were submitted to IMG and subsequently downloaded on July 22, 2022. The number of genes in each pfam were then averaged across all genomes from the order. Gene counts were compared statistically using a Fisher’s exact test (*82*) to determine which enzyme families have a higher gene count in the isolate genome than the average of >1000 genomes from the same bacterial Order. Those pfams having a gene count log base 2-fold change greater than one were selected for further study.

### Gene synthesis, cloning, and enzyme expression

Protein sequences were codon optimized for *E. coli* and purchased from Twist Biosciences (South San Francisco, CA, USA) either in the pET-28a vector, or as gene fragments. Gene fragments were cloned into the pET-28a vector by using restriction-ligation cloning at the BamHI and HindIII restriction enzyme sites. Upon selection of colonies using kanamycin as a resistance marker, plasmids were transformed into *E. coli* BL21(DE3) using standard heat shock transformation. Each construct had a 6x histidine tag on the N or C terminus for purification.

For protein expression, BL21 strains containing the plasmid of interest were inoculated at 37°C until they reached an optical density of approximately 0.7 OD600. 0.3 mM IPTG was added to each culture for induction of protein expression. Cultures were then grown at 30 °C for 6 hours to express protein, which were then visualized via SDS-PAGE. For proteins that did not express at 30 °C (*Cv*DyP, and subsequently all DyPs for the remainder of the study), cultures were instead grown for 18–22 hours at 18 °C.

### Protein purification

After protein expression is complete, cultures were pelleted, lysed using Solulyse bacterial protein extraction reagent (Genlantis Inc. San Diego, CA, USA. Catalog number L100500) per the manufacturer’s instructions. Soluble protein fraction in Solulyse solution was brought to a concentration of 10 mM imidazole and then purified, through nickel affinity chromatography via FPLC. Using a HisTrap HP 1 mL column (Cytiva, Marlborough, MA, USA), binding buffer was comprised of 20 mM NaH_2_PO_4_, 500 mM NaCl, and 20 mM Imidazole and elution buffer was comprised of 20 mM NaH_2_PO_4_, 500 mM NaCl, and 500 mM Imidazole. A gradient from 0–50% elution buffer, followed by isocratic flow of 100% of elution buffer was used to purify the protein. All purifications were carried out at 4°C.

### Production of stripped LDPE films

LDPE (low-density polyethylene pellets [melt index:25 g/10 min] were purchased from Sigma Aldrich Chemical Company, St. Louis, MO, USA; catalog number 428043 – 250 g) (7.2 g) was dissolved in xylenes (150 mL) under reflux (130 °C) with constant stirring for 3 hours. The solution was left to cool to 60 °C without stirring and ice-cold methanol (19.95 mL) was added dropwise before pouring the solution into room temperature methanol (129.2 mL). The solution was vacuum filtered, and the recovered polymer washed with fresh methanol 3 times before drying at room temperature for 48 hours. Recovered LDPE (7–8 g) was added to a cellulose extraction thimble and placed in a Soxhlet apparatus with chloroform (200 mL). The heating rate was controlled to achieve Soxhlet extraction cycles of ∼15 minutes duration over a total of 24 hours. The remaining polymer was left to dry at room temperature for 1 hour and dried under vacuum for 16 hours. The stripped PE was then hot pressed between Kapton films at 11 MPa at 180 °C for 5 minutes to form thin films (< 1 mm).

### Enzyme assaying on plastic films

For activity screening, proteins were heterologously expressed in *E. coli* BL21(DE3) and crude lysates were dosed onto LDPE films (Goodfellow Cambridge Limited, Huntingdon, England. Catalog number LS580568) in three 90-minute doses and allowed to dry overnight prior to screening for oxygenation via FTIR. A more thorough analysis of enzyme activity on films started with purified enzymes that were stored at a concentration of 1.0 g/L, verified by a Bradford assay. To screen enzymes for activity, 10 µL of 1.0 g/L enzyme was dosed onto a film (Goodfellow Cambridge Limited, Huntingdon, England. Catalog number LS580568 or LDPE stripped of additives from (Sigma Aldrich Chemical Company, St. Louis, MO, USA; catalog number 428043 – 250 g) with 1 mM of the appropriate cofactor in a sodium phosphate buffer to a final volume of 30 µL. Three, two-hour doses of this nature were performed, with the last dose allowed to react and dry overnight for 16 hours. Enzyme re-dosing was deemed necessary as a single dose saturating the surface proved insufficient for deconstruction. Moreover, the film surface dried as liquid evaporates, requiring enzyme to be re-dosed to sustain the reaction. Each enzyme was initially tested at pH 7, for ease of assaying. Plastic films were then washed in water and ethanol, air dried, and analyzed via FTIR to monitor chemical changes. Enzymatic reactions of DyPs were carried out at 1 mM H_2_O_2_ (Sigma Aldrich Chemical Company. St. Louis, MO, USA Catalog number H1009 – 500ML) and a pH of 4.0 as a result of enzyme optimization studies (*83–85*).

### Pyrogallol peroxidase assay

Pyrogallol (Sigma Aldrich Chemical Company. St. Louis, MO, USA Catalog number P0381 – 25 g) was used as a standard substrate for measuring enzyme activity of peroxidases. The enzyme assay used followed the ‘Enzymatic Assay of Peroxidase (EC 1.11.1.7)’ protocol from Millipore sigma (*86*). Briefly, 0.027% v/v hydrogen peroxide was added with 0.5% w/v pyrogallol and 0.75 units of peroxidase for the reaction. The generation of purpurgallin was measured using absorbance at 420 nm.

### DyP-Peroxidase sequence similarity screening

From the metagenomic data in ‘Metagenomic data analysis to identify protein families of interest’, 6 enzymes were randomly selected from LDPE fed mealworm gut sample that fell within pfam 04261 and were thus classified as dye-decolorizing peroxidases. Additionally, a protein BLAST search on *Cv*DyP was performed to find enzymes with similar sequences from the same genus (*87*). From this search, two hits within the top 10 BLAST results by E-value were specifically selected to ensure that the test database encompassed the four subclasses of class I DyPs per classification in (*53*, *54*). This list was generated to encompass a variety of DyP subclasses and taxonomic origins, including several in the same I3 subclass that were from the same genus as *Cv*DyP. In total, 9 enzymes across class I DyPs were tested for PE activity in addition to *CvDyP* (Supplementary Table 5). Species with DyPs included in this list were: *Rothia halotolerans (RhDyP), unclassified Brevibacterium (UnBDyp), Brevibacterium oceani (BoDyP), Brevibacterium linens (BlDyP, Bl2DyP, BL3DyP), Corynebacterium provencense (CpDyP), Corynebacterium sp. (CsDyP)* and *Corynebacterium neomassiliense (CnDyP)*; where *CsDyP and CnDyP* are those from the BLAST search and all others listed are from the LDPE-fed metagenomic data.

### DyP-Peroxidase sequence secretion tag identification and structure prediction

All sequences were screened using signalP 6.0 using the fast model (*88*). The fasta file produced with secretion tags cleaved off was used to generate AlphaFold predictions of structure via Alphafold colab (*89–93*). Additionally, an outgroup protein of a similar peroxidase coming from the same genus as that of one of the screened DyP peroxidases was used. The outgroup protein used the AlphaFold prediction available on interpro and the protein has accession number A0A163AWC0, identified as a catalase peroxidase (*94*).

### YRB map generation

YRB maps were generated according to Hagemans et. al. (*60*). The python script from the original manuscript was downloaded and used to generate YRB maps.

### *Cp*-CNHL activity assay on C10-C40 alkanes

750 µL C10-C40 alkane standards (Sigma Aldrich, catalog #68281) were run through a rotary evaporator at 150 mbar absolute pressure, 50°C jacket, -20°c condenser, 200 rpm rotation for 5 minutes and dried in a vacuum oven overnight at 150 mbar absolute pressure at 100°C, left uncapped overnight. pH4 buffer, Cp+CNHL and H_2_O_2_ were applied directly to dried alkane wax emulating “**Enzyme assaying on plastic films.”** Alkanes were resuspended into 150 µL hexane to re-dissolve alkanes for gas chromatography (GC) analysis. 2-decanone (Sigma Aldrich, 196207 catalog #) was added at 1 mg/mL as an internal standard. Chromatographic analyses were performed with a gas chromatography-flame ionization detector system (GC-FID) 7980A-5975C from Agilent Technologies. Separation of the metabolites was performed on a DB-5 Column coated with polyimide (30 m length, 0.25 mm inner diameter, and 0.1 µm film thickness; Agilent Technologies, USA) for proper separation of substances, and Helium (He) was utilized as a carrier gas. The analysis was performed using a split injector at 350 °C and an injection volume of 1 µl. The ion source temperature was 230 °C, mass spectral analysis was performed in scan mode, the quadrupole temperature of 150 °C, and a fragmentation voltage of 70 eV.

### Cp-CNHL molecular docking analysis

Molecular docking simulations were prepared using Autodock 4 in Dockey for MacOS version Big Sur 11.7. A custom bounding box was drawn around the hydrophobic loop of the enzyme when docking dotriacontane and a custom bounding box was drawn around the aromatic residues W300 and Y424 when docking reactive blue. A custom bounding box was drawn around the active site when docking decane and dotriacontane to the active site of the enzyme. The Lamarckian GA property search algorithm, using Autodock 4 default parameters. Ligands were prepared using default parameters on prepare_ligand4 ligand preparation tool. Ligands were ZINC6920423 (dotriacontane), CHEMBL134537 (decane), and CHEMBL5187239.

## Acknowledgements

This research is supported by the U.S. Department of Energy, Office of Science, Office of Biological and Environmental Research under Award Numbers DE-SC0022018 and DE-SC0023085. This research is supported as part of the Center for Plastics Innovation, an Energy Frontier Research Center funded by the U.S. Department of Energy (DOE), Office of Science, Basic Energy Sciences (BES), under award DE-SC0021166 (HT-SEC and substrate production). RRK was funded in part by the Delaware Environmental Institute, University of Delaware and the Chemistry Biology Interface at the University of Delaware, under NIH training grant T32GM133395. DAH was funded in part by the U.S. Department of Education GAANN (P200A210065). Support from the University of Delaware CBCB Bioinformatics Data Science Core Facility (RRID:SCR_017696) including use of the BIOMIX and BioStore computational resources was made possible through funding from Delaware INBRE (NIGMS P20GM103446), NIH Shared Instrumentation Grant (S10OD028725) the State of Delaware, and the Delaware Biotechnology Institute. Reagents for this research were ordered in part with funds from a QIAGEN Young Scientist Research Grant award to RRK.

We thank the Advanced Materials Characterization lab at the University of Delaware for FTIR instrument time. XPS analysis was performed with the instrument sponsored by the National Science Foundation under grant No. CHE-1428149.

A portion of this research was performed on a project award (doi: 10.46936/fics.proj.2021.60038/60000396) under the FICUS program and used resources at the DOE Joint Genome Institute and the Environmental Molecular Sciences Laboratory, which are DOE Office of Science User Facilities. Both facilities are sponsored by the Biological and Environmental Research program and operated under Contract Nos. DE-AC02-05CH11231 (JGI) and DE-AC05-76RL01830 (EMSL). The work conducted by the U.S. Department of Energy Joint Genome Institute (https://ror.org/04xm1d337), a DOE Office of Science User Facility, is supported by the Office of Science of the U.S. Department of Energy operated under Contract No. DE-AC02-05CH11231.” We would like to thank the following personnel at the Joint Genome Institute for their efforts in genome sequencing: Marcel Huntemann, Kurt LaButti, Alex Spunde, Krishnaveni Palaniappan, Stephan Ritter, Natalia Mikhailova, I-Min A. Chen, Dimitrios Stamatis, T.B.K. Reddy, Chris Daum, Miranda Harmon-Smith, Natalia Ivanova, Nikos C. Kyrpides, and Tanja Woyke

## Author Contributions

Ross R. Klauer: led all experimental work, investigation, methodology, validation, formal analysis, writing – original draft

Lummy Oliveira Monteiro: investigation

D. Alex Hansen: data curation, software, assisted in writing original draft

Zoé O. G. Schyns: investigation, formal analysis, reviewed & edited the manuscript

Jenna A. Moore-Ott: assisted in formal analysis of data

Mekhi Williams: investigation

Megan Tarr: investigation

Jyoti Singh: investigation

Ashwin Mhadeshwar: investigation

LaShanda T. J. Korley: reviewed and edited the manuscript

Kevin Solomon and Mark Blenner: conceptualized the study, supervised the study, acquired funding, and reviewed & edited the manuscript.

The order of the two co-corresponding authors was determined by a best of three odds and evens match between KVS and MAB.

## References

1. Global Plastics Outlook: Plastic waste in 2019 (Edition 2022), OECD (2017); 10.1787/9b6fb836-en.

2. H. Li, H. A. Aguirre-Villegas, R. D. Allen, X. Bai, C. H. Benson, G. T. Beckham, S. L. Bradshaw, J. L. Brown, R. C. Brown, V. S. Cecon, J. B. Curley, G. W. Curtzwiler, S. Dong, S. Gaddameedi, J. E. García, I. Hermans, M. S. Kim, J. Ma, L. O. Mark, M. Mavrikakis, O. O. Olafasakin, T. A. Osswald, K. G. Papanikolaou, H. Radhakrishnan, M. A. Sanchez Castillo, K. L. Sánchez-Rivera, K. N. Tumu, R. C. Van Lehn, K. L. Vorst, M. M. Wright, J. Wu, V. M. Zavala, P. Zhou, G. W. Huber, Expanding plastics recycling technologies: chemical aspects, technology status and challenges. Green Chemistry 24, 8899–9002 (2022).

3. L. T. J. Korley, T. H. Epps, B. A. Helms, A. J. Ryan, Toward polymer upcycling—adding value and tackling circularity. Science 373, 66–69 (2021).

4. R. Geyer, J. R. Jambeck, K. L. Law, Production, use, and fate of all plastics ever made. Science Advances 3 (2017).

5. Z. R. Hinton, M. R. Talley, P. A. Kots, A. V. Le, T. Zhang, M. E. Mackay, A. M. Kunjapur, P. Bai, D. G. Vlachos, M. P. Watson, M. C. Berg, Thomas H. Epps, L. T. J. Korley, Innovations Toward the Valorization of Plastics Waste. Annual Review of Materials Research 52, 249–280 (2022).

6. G. Lopez, M. Artetxe, M. Amutio, J. Bilbao, M. Olazar, Thermochemical routes for the valorization of waste polyolefinic plastics to produce fuels and chemicals. A review. Renewable and Sustainable Energy Reviews 73, 346–368 (2017).

7. A. Rahimi, J. M. García, Chemical recycling of waste plastics for new materials production. Nature Reviews Chemistry 1 (2017).

8. L. D. Ellis, N. A. Rorrer, K. P. Sullivan, M. Otto, J. E. McGeehan, Y. Román-Leshkov, N. Wierckx, G. T. Beckham, Chemical and biological catalysis for plastics recycling and upcycling. Nat Catal 4, 539–556 (2021).

9. R. R. Klauer, D. A. Hansen, D. Wu, L. M. O. Monteiro, K. V. Solomon, M. A. Blenner, Biological Upcycling of Plastics Waste. Annu Rev Chem Biomol Eng, doi: 10.1146/annurev-chembioeng-100522-115850 (2024).

10. H. Lu, D. J. Diaz, N. J. Czarnecki, C. Zhu, W. Kim, R. Shroff, D. J. Acosta, B. R. Alexander, H. O. Cole, Y. Zhang, N. A. Lynd, A. D. Ellington, H. S. Alper, Machine learning-aided engineering of hydrolases for PET depolymerization. Nature 2022 604:7907 604, 662–667 (2022).

11. H. F. Son, I. J. Cho, S. Joo, H. Seo, H.-Y. Sagong, S. Y. Choi, S. Y. Lee, K.-J. Kim, Rational Protein Engineering of Thermo-Stable PETase from Ideonella sakaiensis for Highly Efficient PET Degradation. ACS Catal. 9, 3519–3526 (2019).

12. Y. Cui, Y. Chen, X. Liu, S. Dong, Y. Tian, Y. Qiao, R. Mitra, J. Han, C. Li, X. Han, W. Liu, Q. Chen, W. Wei, X. Wang, W. Du, S. Tang, H. Xiang, H. Liu, Y. Liang, K. N. Houk, B. Wu, Computational Redesign of a PETase for Plastic Biodegradation under Ambient Condition by the GRAPE Strategy. ACS Catal. 11, 1340–1350 (2021).

13. V. Tournier, C. M. Topham, A. Gilles, B. David, C. Folgoas, E. Moya-Leclair, E. Kamionka, M.-L. Desrousseaux, H. Texier, S. Gavalda, M. Cot, E. Guémard, M. Dalibey, J. Nomme, G. Cioci, S. Barbe, M. Chateau, I. André, S. Duquesne, A. Marty, An engineered PET depolymerase to break down and recycle plastic bottles. Nature 580, 216–219 (2020).

14. Y. Cui, Y. Chen, J. Sun, T. Zhu, H. Pang, C. Li, W.-C. Geng, B. Wu, Computational redesign of a hydrolase for nearly complete PET depolymerization at industrially relevant high-solids loading. Nat Commun 15, 1417 (2024).

15. N. Mohanan, Z. Montazer, P. K. Sharma, D. B. Levin, Microbial and Enzymatic Degradation of Synthetic Plastics. Frontiers in Microbiology 11 (2020).

16. A. Sanluis-Verdes, P. Colomer-Vidal, F. Rodriguez-Ventura, M. Bello-Villarino, M. Spinola-Amilibia, E. Ruiz-Lopez, R. Illanes-Vicioso, P. Castroviejo, R. A. Cigliano, M. Montoya, P. Falabella, C. Pesquera, L. Gonzalez-Legarreta, E. Arias-Palomo, M. Solà, T. Torroba, C. F. Arias, F. Bertocchini, Wax worm saliva and the enzymes therein are the key to polyethylene degradation by Galleria mellonella. doi: 10.1038/s41467-022-33127-w.

17. M. Santo, R. Weitsman, A. Sivan, The role of the copper-binding enzyme – laccase – in the biodegradation of polyethylene by the actinomycete Rhodococcus ruber. International Biodeterioration & Biodegradation 84, 204–210 (2013).

18. A. A. Stepnov, E. Lopez-Tavera, R. Klauer, C. L. Lincoln, R. R. Chowreddy, G. T. Beckham, V. G. H. Eijsink, K. Solomon, M. Blenner, G. Vaaje-Kolstad, Revisiting the activity of two poly(vinyl chloride)- and polyethylene-degrading enzymes. bioRxiv [Preprint] (2024). 10.1101/2024.03.15.585159.

19. R. Wei, T. Tiso, J. Bertling, K. O’Connor, L. M. Blank, U. T. Bornscheuer, Possibilities and limitations of biotechnological plastic degradation and recycling. Nat Catal 3, 867–871 (2020).

20. A. M. Brandon, S. H. Gao, R. Tian, D. Ning, S. S. Yang, J. Zhou, W. M. Wu, C. S. Criddle, Biodegradation of Polyethylene and Plastic Mixtures in Mealworms (Larvae of Tenebrio molitor) and Effects on the Gut Microbiome. Environmental science & technology 52, 6526–6533 (2018).

21. L. Jin, P. Feng, Z. Cheng, D. Wang, Effect of biodegrading polyethylene, polystyrene, and polyvinyl chloride on the growth and development of yellow mealworm (Tenebrio molitor) larvae. Environmental Science and Pollution Research 30, 37118–37126 (2022).

22. S.-S. Yang, M.-Q. Ding, L. He, C.-H. Zhang, Q.-X. Li, D.-F. Xing, G.-L. Cao, L. Zhao, J. Ding, N.-Q. Ren, W.-M. Wu, Biodegradation of polypropylene by yellow mealworms (Tenebrio molitor) and superworms (Zophobas atratus) via gut-microbe-dependent depolymerization. Science of The Total Environment 756, 144087 (2021).

23. B.-Y. Peng, Y. Su, Z. Chen, J. Chen, X. Zhou, M. E. Benbow, C. S. Criddle, W.-M. Wu, Y. Zhang, Biodegradation of Polystyrene by Dark (Tenebrio obscurus) and Yellow (Tenebrio molitor) Mealworms (Coleoptera: Tenebrionidae). Environ. Sci. Technol. 53, 5256–5265 (2019).

24. S.-S. Yang, A. M. Brandon, J. C. Andrew Flanagan, J. Yang, D. Ning, S.-Y. Cai, H.-Q. Fan, Z.-Y. Wang, J. Ren, E. Benbow, N.-Q. Ren, R. M. Waymouth, J. Zhou, C. S. Criddle, W.-M. Wu, Biodegradation of polystyrene wastes in yellow mealworms (larvae of Tenebrio molitor Linnaeus): Factors affecting biodegradation rates and the ability of polystyrene-fed larvae to complete their life cycle. Chemosphere 191, 979–989 (2018).

25. A. Chamas, H. Moon, J. Zheng, Y. Qiu, T. Tabassum, J. H. Jang, M. Abu-Omar, S. L. Scott, S. Suh, Degradation Rates of Plastics in the Environment. ACS Sustainable Chem. Eng. 8, 3494–3511 (2020).

26. H. R. Kim, C. Lee, H. Shin, H. Y. Koh, S. Lee, D. Choi, Interplay Between Superworm and its Gut Microbiome in Facilitating Polyethylene Biodegradation by Host Transcriptomic Analysis: Insights from Xenobiotic Metabolism. doi: 10.21203/rs.3.rs-2815027/v1 (2023).

27. S. Jiang, T. Su, J. Zhao, Z. Wang, Biodegradation of Polystyrene by Tenebrio molitor, Galleria mellonella, and Zophobas atratus Larvae and Comparison of Their Degradation Effects. Polymers 13, 3539 (2021).

28. Y. Yang, J. Yang, W. M. Wu, J. Zhao, Y. Song, L. Gao, R. Yang, L. Jiang, Biodegradation and Mineralization of Polystyrene by Plastic-Eating Mealworms: Part 1. Chemical and Physical Characterization and Isotopic Tests. Environmental Science and Technology 49, 12080–12086 (2015).

29. Y. Yang, J. Yang, W.-M. Wu, J. Zhao, Y. Song, L. Gao, R. Yang, L. Jiang, Biodegradation and Mineralization of Polystyrene by Plastic-Eating Mealworms: Part 2. Role of Gut Microorganisms. Environ. Sci. Technol. 49, 12087–12093 (2015).

30. S. W. Przemieniecki, A. Kosewska, S. Ciesielski, O. Kosewska, Changes in the gut microbiome and enzymatic profile of Tenebrio molitor larvae biodegrading cellulose, polyethylene and polystyrene waste. Environmental Pollution 256, 113265 (2020).

31. B. J. Cassone, C. M. R. Lemoine, H. C. Grove, O. Elebute, S. M. P. Villanueva, Role of the intestinal microbiome in low-density polyethylene degradation by caterpillar larvae of the greater wax moth, Galleria mellonella. doi: 10.1098/rspb.2020.0112 (2020).

32. Y. P. Khanna, E. A. Turi, T. J. Taylor, V. V. Vickroy, R. F. Abbott, Dynamic mechanical relaxations in polyethylene. Macromolecules 18, 1302–1309 (1985).

33. R. B. Richards, Polyethylene-structure, crystallinity and properties. Journal of Applied Chemistry 1, 370–376 (1951).

34. Y. Tokiwa, B. P. Calabia, C. U. Ugwu, S. Aiba, Biodegradability of Plastics. International Journal of Molecular Sciences 10, 3722–3742 (2009).

35. S.-S. Yang, M.-Q. Ding, Z.-R. Zhang, J. Ding, S.-W. Bai, G.-L. Cao, L. Zhao, J.-W. Pang, D.-F. Xing, N.-Q. Ren, W.-M. Wu, Confirmation of biodegradation of low-density polyethylene in dark-versus yellow-mealworms (larvae of *Tenebrio obscurus* versus *Tenebrio molitor*) via. gut microbe-independent depolymerization. Science of The Total Environment 789, 147915 (2021).

36. Q. Wang, H. Chen, W. Gu, S. Wang, Y. Li, Biodegradation of aged polyethylene (PE) and polystyrene (PS) microplastics by yellow mealworms (*Tenebrio molitor* larvae). Science of The Total Environment 927, 172243 (2024).

37. J. Yang, Y. Yang, W.-M. Wu, J. Zhao, L. Jiang, Evidence of Polyethylene Biodegradation by Bacterial Strains from the Guts of Plastic-Eating Waxworms. Environ. Sci. Technol. 48, 13776–13784 (2014).

38. H. Lou, R. Fu, T. Long, B. Fan, C. Guo, L. Li, J. Zhang, G. Zhang, Biodegradation of polyethylene by Meyerozyma guilliermondii and Serratia marcescens isolated from the gut of waxworms (larvae of Plodia interpunctella). Science of The Total Environment 853, 158604 (2022).

39. F. Rojo, Degradation of alkanes by bacteria. Environmental Microbiology 11, 2477–2490 (2009).

40. Y. Ji, G. Mao, Y. Wang, M. Bartlam, Structural insights into diversity and n-alkane biodegradation mechanisms of alkane hydroxylases. Frontiers in Microbiology 4, 58–58 (2013).

41. J. B. Van Beilen, Z. Li, W. A. Duetz, T. H. M. Smits, B. Witholt, Diversity of Alkane Hydroxylase Systems in the Environment. Oil & Gas Science and Technology 58, 427–440 (2003).

42. T. L. (Tom L. Cottrell, The strengths of chemical bonds. Butterworths Scienticif, Academic Press edition, 290–308 (1954).

43. S. Greer, M. Wen, D. Bird, X. Wu, L. Samuels, L. Kunst, R. Jetter, The Cytochrome P450 Enzyme CYP96A15 Is the Midchain Alkane Hydroxylase Responsible for Formation of Secondary Alcohols and Ketones in Stem Cuticular Wax of Arabidopsis. Plant Physiol 145, 653–667 (2007).

44. A. Gutiérrez, E. D. Babot, R. Ullrich, M. Hofrichter, A. T. Martínez, J. C. del Río, Regioselective oxygenation of fatty acids, fatty alcohols and other aliphatic compounds by a basidiomycete heme-thiolate peroxidase. Archives of Biochemistry and Biophysics 514, 33–43 (2011).

45. Q. Wu, B. Qu, Y. Xu, Q. Wu, Surface photo-oxidation and photostabilization of photocross-linked polyethylene. Polymer Degradation and Stability 68, 97–102 (2000).

46. M. R. Sanchis, V. Blanes, M. Blanes, D. Garcia, R. Balart, Surface modification of low density polyethylene (LDPE) film by low pressure O2 plasma treatment. European Polymer Journal 42, 1558–1568 (2006).

47. A. M. Deschamps, G. Mahoudeau, J. M. Lebeault, Fast degradation of kraft lignin by bacteria. European J. Appl. Microbiol. Biotechnol. 9, 45–51 (1980).

48. S. Baghel, J. Anandkumar, Biodepolymerization of Kraft lignin for production and optimization of vanillin using mixed bacterial culture. Bioresource Technology Reports 8, 100335 (2019).

49. P. Boruah, P. Sarmah, P. K. Das, T. Goswami, Exploring the lignolytic potential of a new laccase producing strain *Kocuria* sp. PBS-1 and its application in bamboo pulp bleaching. International Biodeterioration & Biodegradation 143, 104726 (2019).

50. A. Bordiean, M. Krzyżaniak, M. J. Stolarski, D. Peni, Growth Potential of Yellow Mealworm Reared on Industrial Residues. Agriculture 10, 599 (2020).

51. C.-L. Yeh, M. A. L. Nikolić, B. Gomes, E. Gauthier, B. Laycock, P. Halley, S. E. Bottle, J. M. Colwell, The effect of common agrichemicals on the environmental stability of polyethylene films. Polymer Degradation and Stability 120, 53–60 (2015).

52. B. Oppert, A. T. Dossey, F.-C. Chu, E. Šatović-Vukšić, M. Plohl, T. P. L. Smith, S. Koren, M. L. Olmstead, D. Leierer, G. Ragan, J. S. Johnston, The Genome of the Yellow Mealworm, Tenebrio molitor: It’s Bigger Than You Think. Genes 14, 2209 (2023).

53. T. Yoshida, Y. Sugano, Unexpected diversity of dye-decolorizing peroxidases. Biochemistry and Biophysics Reports 33, 101401 (2023).

54. T. Yoshida, Y. Sugano, A structural and functional perspective of DyP-type peroxidase family. Archives of Biochemistry and Biophysics 574, 49–55 (2015).

55. D. Maizel, S. M. Utturkar, S. D. Brown, M. A. Ferrero, B. P. Rosen, Draft Genome Sequence of Brevibacterium linens AE038-8, an Extremely Arsenic-Resistant Bacterium. Genome Announcements 3 (2015).

56. E. Alexa, J. F. Cobo-Diaz, E. Renes, T. F. ÓCallaghan, K. Kilcawley, D. Mannion, I. Skibinska, L. Ruiz, A. Margolles, P. Fernández-Gómez, A. Alvarez-Molina, F. Crispie, P. Puente-Gómez, M. López, M. Prieto, P. Cotter, A. Alvarez-Ordóñez, Environmental sources along natural cave ripening shape the microbiome and metabolome of artisanal blue-veined cheeses. [Preprint] (2021). 10.21203/rs.3.rs-1150288/v1.

57. S. Ndongo, C. Andrieu, P.-E. Fournier, J.-C. Lagier, D. Raoult, ‘Actinomyces provencensis’ sp. nov., ‘Corynebacterium bouchesdurhonense’ sp. nov., ‘Corynebacterium provencense’ sp. nov. and ‘Xanthomonas massiliensis’ sp. nov., 4 new species isolated from fresh stools of obese French patients. New Microbes and New Infections 18, 24 (2017).

58. M. Boxberger, I. Hasni, M. Bilen, B. La Scola, *Corynebacterium neomassiliense* sp. nov., a new bacterium isolated in a stool sample from a healthy male pygmy. New Microbes and New Infections 34, 100644 (2020).

59. J. Jumper, R. Evans, A. Pritzel, T. Green, M. Figurnov, O. Ronneberger, K. Tunyasuvunakool, R. Bates, A. Žídek, A. Potapenko, A. Bridgland, C. Meyer, S. A. A. Kohl, A. J. Ballard, A. Cowie, B. Romera-Paredes, S. Nikolov, R. Jain, J. Adler, T. Back, S. Petersen, D. Reiman, E. Clancy, M. Zielinski, M. Steinegger, M. Pacholska, T. Berghammer, S. Bodenstein, D. Silver, O. Vinyals, A. W. Senior, K. Kavukcuoglu, P. Kohli, D. Hassabis, Highly accurate protein structure prediction with AlphaFold. Nature 596, 583– 589 (2021).

60. D. Hagemans, I. A. E. M. van Belzen, T. Morán Luengo, S. G. D. Rüdiger, A script to highlight hydrophobicity and charge on protein surfaces. Front Mol Biosci 2, 56 (2015).

61. R. Shrestha, G. Huang, D. A. Meekins, B. V. Geisbrecht, P. Li, Mechanistic Insights into Dye-Decolorizing Peroxidase Revealed by Solvent Isotope and Viscosity Effects. ACS Catal 7, 6352–6364 (2017).

62. D. Linde, R. Pogni, M. Cañellas, F. Lucas, V. Guallar, M. C. Baratto, A. Sinicropi, V. Sáez-Jiménez, C. Coscolín, A. Romero, F. J. Medrano, F. J. Ruiz-Dueñas, A. T. Martínez, Catalytic surface radical in dye-decolorizing peroxidase: a computational, spectroscopic and site-directed mutagenesis study. Biochem J 466, 253–262 (2015).

63. R. Shrestha, X. Chen, K. X. Ramyar, Z. Hayati, E. A. Carlson, S. H. Bossmann, L. Song, B. V. Geisbrecht, P. Li, Identification of Surface-Exposed Protein Radicals and A Substrate Oxidation Site in A-Class Dye-Decolorizing Peroxidase from Thermomonospora curvata. ACS Catal. 6, 8036–8047 (2016).

64. A. K. Chaudhary, R. P. Vijayakumar, Effect of chemical treatment on biological degradation of high-density polyethylene (HDPE). Environ Dev Sustain 22, 1093–1104 (2020).

65. J. L. Brown, E. Rodriguez-Ocasio, C. Peterson, M. Blenner, R. Smith, L. Jarboe, R. Brown, High-Temperature, Noncatalytic Oxidation of Polyethylene to a Fermentation Substrate Robustly Utilized by Candida maltosa. ACS Sustainable Chem. Eng. 11, 17778–17786 (2023).

66. C. Yao, W. Xia, M. Dou, Y. Du, J. Wu, Oxidative degradation of UV-irradiated polyethylene by laccase-mediator system. Journal of Hazardous Materials 440, 129709–129709 (2022).

67. S. Mukherjee, P. P. Kundu, Alkaline fungal degradation of oxidized polyethylene in black liquor: Studies on the effect of lignin peroxidases and manganese peroxidases. Journal of Applied Polymer Science 131 (2014).

68. H. J. Jeon, M. N. Kim, Functional analysis of alkane hydroxylase system derived from Pseudomonas aeruginosa E7 for low molecular weight polyethylene biodegradation. International Biodeterioration and Biodegradation 103, 141–146 (2015).

69. J. Zampolli, M. Mangiagalli, D. Vezzini, M. Lasagni, D. Ami, A. Natalello, F. Arrigoni, L. Bertini, M. Lotti, P. Di Gennaro, Oxidative degradation of polyethylene by two novel laccase-like multicopper oxidases from *Rhodococcus opacus* R7. Environmental Technology & Innovation 32, 103273 (2023).

70. M. Ahmad, J. N. Roberts, E. M. Hardiman, R. Singh, L. D. Eltis, T. D. H. Bugg, Identification of DypB from Rhodococcus jostii RHA1 as a Lignin Peroxidase. Biochemistry 50, 5096– 5107 (2011).

71. C. Yang, F. Yue, Y. Cui, Y. Xu, Y. Shan, B. Liu, Y. Zhou, X. Lü, Biodegradation of lignin by Pseudomonas sp. Q18 and the characterization of a novel bacterial DyP-type peroxidase. Journal of Industrial Microbiology and Biotechnology 45, 913–927 (2018).

72. R. Rahmanpour, T. D. H. Bugg, Characterisation of Dyp-type peroxidases from *Pseudomonas fluorescens* Pf-5: Oxidation of Mn(II) and polymeric lignin by Dyp1B. Archives of Biochemistry and Biophysics 574, 93–98 (2015).

73. Y. Du, C. Yao, M. Dou, J. Wu, L. Su, W. Xia, Oxidative degradation of pre-oxidated polystyrene plastics by dye decolorizing peroxidases from Thermomonospora curvata and Nostocaceae. Journal of Hazardous Materials 436, 129265–129265 (2022).

74. K. Nys, P. G. Furtmüller, C. Obinger, S. Van Doorslaer, V. Pfanzagl, On the Track of Long-Range Electron Transfer in B-Type Dye-Decolorizing Peroxidases: Identification of a Tyrosyl Radical by Computational Prediction and Electron Paramagnetic Resonance Spectroscopy. Biochemistry 60, 1226–1241 (2021).

75. Z. R. Hinton, P. A. Kots, M. Soukaseum, B. C. Vance, D. G. Vlachos, T. H. Epps, L. T. J. Korley, Antioxidant-induced transformations of a metal-acid hydrocracking catalyst in the deconstruction of polyethylene waste. Green Chemistry 24, 7332–7339 (2022).

76. Blood Agar Plates and Hemolysis, ASM.org. https://asm.org:443/Protocols/Blood-Agar-Plates-and-Hemolysis-Protocols.

77. Polyethylene Surfaces. http://www.xpsfitting.com/2015/11/polyethylene-surfaces.html.

78. S. Bennett, Solexa Ltd. Pharmacogenomics 5, 433–438 (2004).

79. BBTools, *DOE* Joint Genome Institute. https://jgi.doe.gov/data-and-tools/software-tools/bbtools/.

80. A. Bankevich, S. Nurk, D. Antipov, A. A. Gurevich, M. Dvorkin, A. S. Kulikov, V. M. Lesin, S. I. Nikolenko, S. Pham, A. D. Prjibelski, A. V. Pyshkin, A. V. Sirotkin, N. Vyahhi, G. Tesler, M. A. Alekseyev, P. A. Pevzner, SPAdes: A New Genome Assembly Algorithm and Its Applications to Single-Cell Sequencing. J Comput Biol 19, 455–477 (2012).

81. I.-M. A. Chen, K. Chu, K. Palaniappan, A. Ratner, J. Huang, M. Huntemann, P. Hajek, S. J. Ritter, C. Webb, D. Wu, N. J. Varghese, T. B. K. Reddy, S. Mukherjee, G. Ovchinnikova, M. Nolan, R. Seshadri, S. Roux, A. Visel, T. Woyke, E. A. Eloe-Fadrosh, N. C. Kyrpides, N. N. Ivanova, The IMG/M data management and analysis system v.7: content updates and new features. Nucleic Acids Research 51, D723–D732 (2023).

82. G. J. G. Upton, Fisher’s Exact Test. Journal of the Royal Statistical Society: Series A (Statistics in Society) 155, 395–402 (1992).

83. T. Uchida, M. Sasaki, Y. Tanaka, K. Ishimori, A Dye-Decolorizing Peroxidase from Vibrio cholerae. Biochemistry 54, 6610–6621 (2015).

84. P. Dhankhar, V. Dalal, J. K. Mahto, B. R. Gurjar, S. Tomar, A. K. Sharma, P. Kumar, Characterization of dye-decolorizing peroxidase from *Bacillus subtilis*. Archives of Biochemistry and Biophysics 693, 108590 (2020).

85. A. Santos, S. Mendes, V. Brissos, L. O. Martins, New dye-decolorizing peroxidases from Bacillus subtilis and Pseudomonas putida MET94: towards biotechnological applications. Appl Microbiol Biotechnol 98, 2053–2065 (2014).

86. Enzymatic Assay of Peroxidase (EC 1.11.1.7). https://www.sigmaaldrich.com/US/en/technical-documents/protocol/protein-biology/enzyme-activity-assays/enzymatic-assay-of-peroxidase.

87. E. W. Sayers, E. E. Bolton, J. R. Brister, K. Canese, J. Chan, D. C. Comeau, R. Connor, K. Funk, C. Kelly, S. Kim, T. Madej, A. Marchler-Bauer, C. Lanczycki, S. Lathrop, Z. Lu, F. Thibaud-Nissen, T. Murphy, L. Phan, Y. Skripchenko, T. Tse, J. Wang, R. Williams, B. W. Trawick, K. D. Pruitt, S. T. Sherry, Database resources of the national center for biotechnology information. Nucleic Acids Res 50, D20–D26 (2022).

88. F. Teufel, J. J. Almagro Armenteros, A. R. Johansen, M. H. Gíslason, S. I. Pihl, K. D. Tsirigos, O. Winther, S. Brunak, G. von Heijne, H. Nielsen, SignalP 6.0 predicts all five types of signal peptides using protein language models. Nat Biotechnol 40, 1023–1025 (2022).

89. M. Mirdita, L. von den Driesch, C. Galiez, M. J. Martin, J. Söding, M. Steinegger, Uniclust databases of clustered and deeply annotated protein sequences and alignments. Nucleic Acids Res. 45, D170–D176 (2017).

90. P. Eastman, J. Swails, J. D. Chodera, R. T. McGibbon, Y. Zhao, K. A. Beauchamp, L.-P. Wang, A. C. Simmonett, M. P. Harrigan, C. D. Stern, R. P. Wiewiora, B. R. Brooks, V. S. Pande, OpenMM 7: Rapid development of high performance algorithms for molecular dynamics. PLOS Comput. Biol. 13 (2017).

91. M. Mirdita, M. Steinegger, J. Söding, MMseqs2 desktop and local web server app for fast, interactive sequence searches. Bioinformatics 35, 2856–2858 (2019).

92. A. L. Mitchell, A. Almeida, M. Beracochea, M. Boland, J. Burgin, G. Cochrane, M. R. Crusoe, V. Kale, S. C. Potter, L. J. Richardson, E. Sakharova, M. Scheremetjew, A. Korobeynikov, A. Shlemov, O. Kunyavskaya, A. Lapidus, R. D. Finn, MGnify: the microbiome analysis resource in 2020. Nucleic Acids Res., doi: 10.1093/nar/gkz1035 (2019).

93. M. Mirdita, K. Schütze, Y. Moriwaki, L. Heo, S. Ovchinnikov, M. Steinegger, ColabFold: making protein folding accessible to all. Nat Methods 19, 679–682 (2022).

94. T. Paysan-Lafosse, M. Blum, S. Chuguransky, T. Grego, B. L. Pinto, G. A. Salazar, M. L. Bileschi, P. Bork, A. Bridge, L. Colwell, J. Gough, D. H. Haft, I. Letunić, A. Marchler-Bauer, H. Mi, D. A. Natale, C. A. Orengo, A. P. Pandurangan, C. Rivoire, C. J. A. Sigrist, I. Sillitoe, N. Thanki, P. D. Thomas, S. C. E. Tosatto, C. H. Wu, A. Bateman, InterPro in 2022. Nucleic Acids Research 51, D418–D427 (2023).

