## Supplementary information file for "Biological polyethylene deconstruction initiated by oxidation from DyP peroxidases"

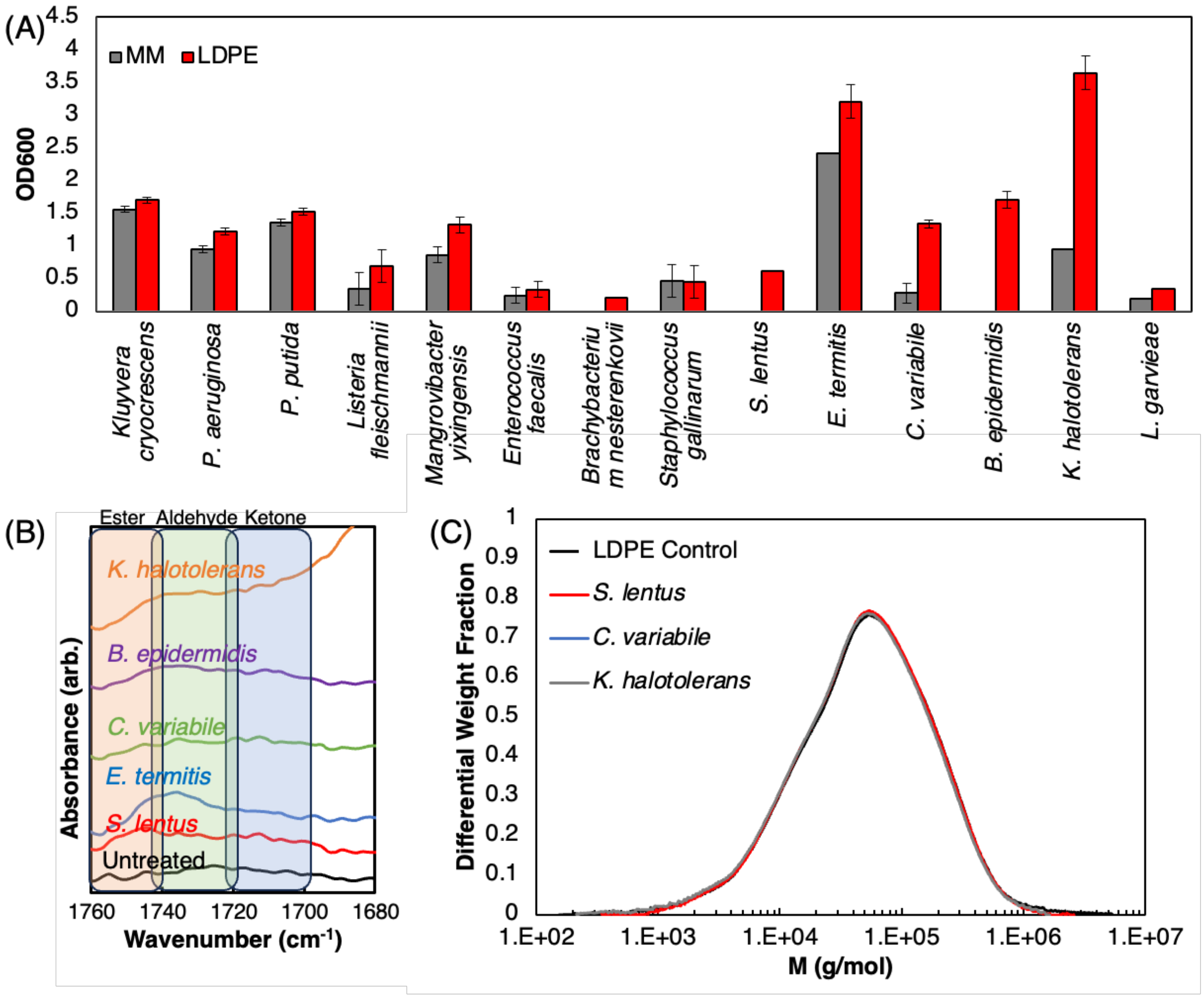


**Supplementary Figure 1: Characterization of yellow mealworm gut microbiota isolates via growth and degradation of PE substrates** (A) Growth data of bacterial isolates in a mineral medium (MM), MM + LDPE powder, or MM + ultra-high molecular weight PE (UHMWPE) powder. OD600 values were recorded after seven days of growth. (B) Zoom in of the carbonyl region in Fig. 1C detailing ranges where aldehydes, ketones, and esters reside in the carbonyl region. (C) SEC chromatograms detailing no deconstruction by select microbial isolates.

**Pseudomonas aeruginosa* and *Pseudomonas putida* were used as control strains and were not isolated from the gut of the yellow mealworm. Full names for each strain can be found in Supplementary Table 1.

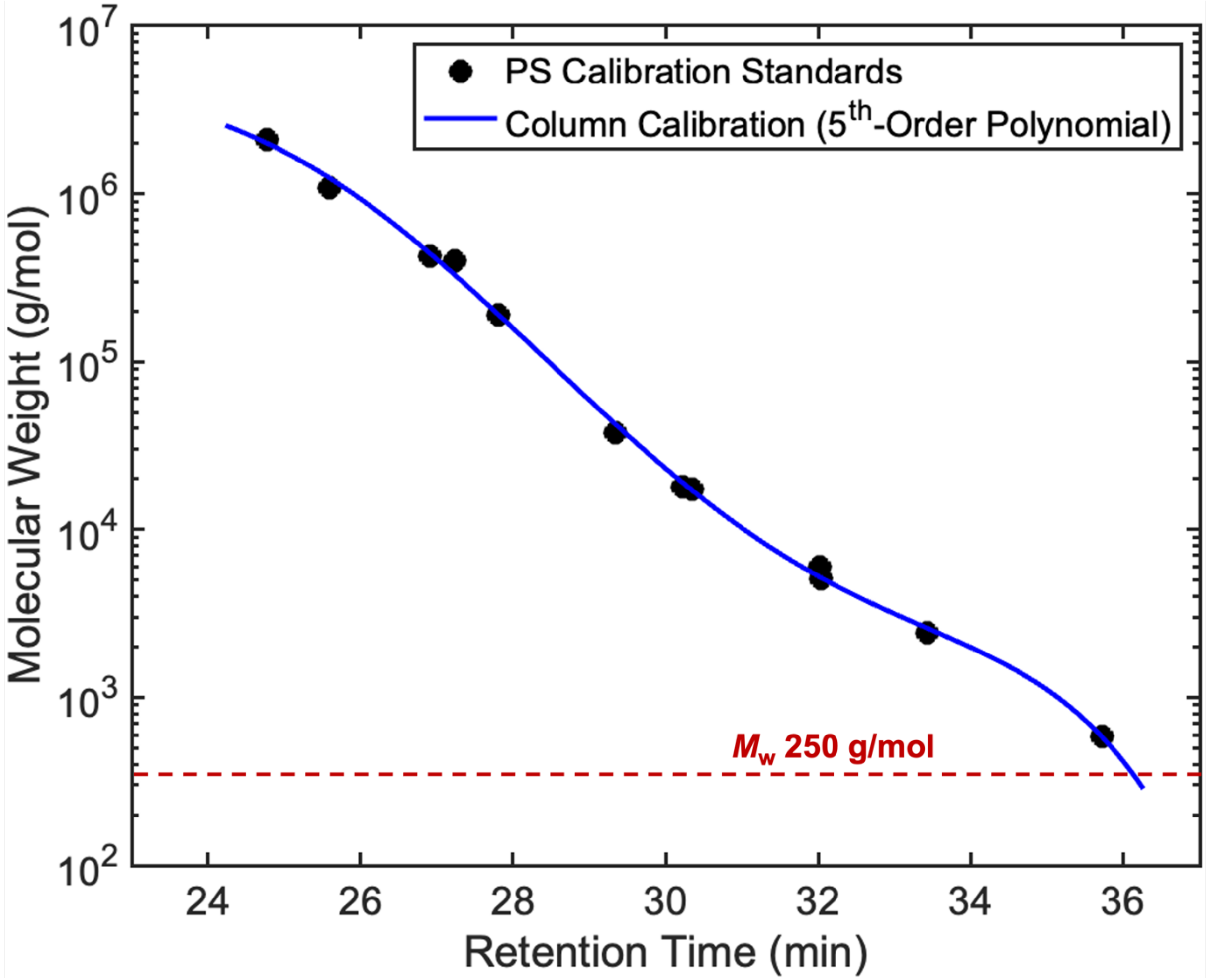


**Supplementary Figure 2 – Column calibration of the HT-GPC instrument**. Calibration was performed from the RI detector using 12 narrow polystyrene standards (6x$10^2$–2x$10^6$ g/mol) at 140 °C in 1,2,4-trichlorobenzene. 5$^{th}$-order calibration curve ($R^2$ > 0.99) calculated using MATLAB's curve fitting toolbox. Due to the molecular weight range chosen for calibration and the column pore size, the upper limit of this 5$^{th}$-order polynomial calibration curve occurs at ~ 36 minutes, which corresponds to ~ 250–300 g/mol. Beyond this retention time, solvent and molecules elute with minimal separation, and thus molecular weights cannot be calculated reliably.

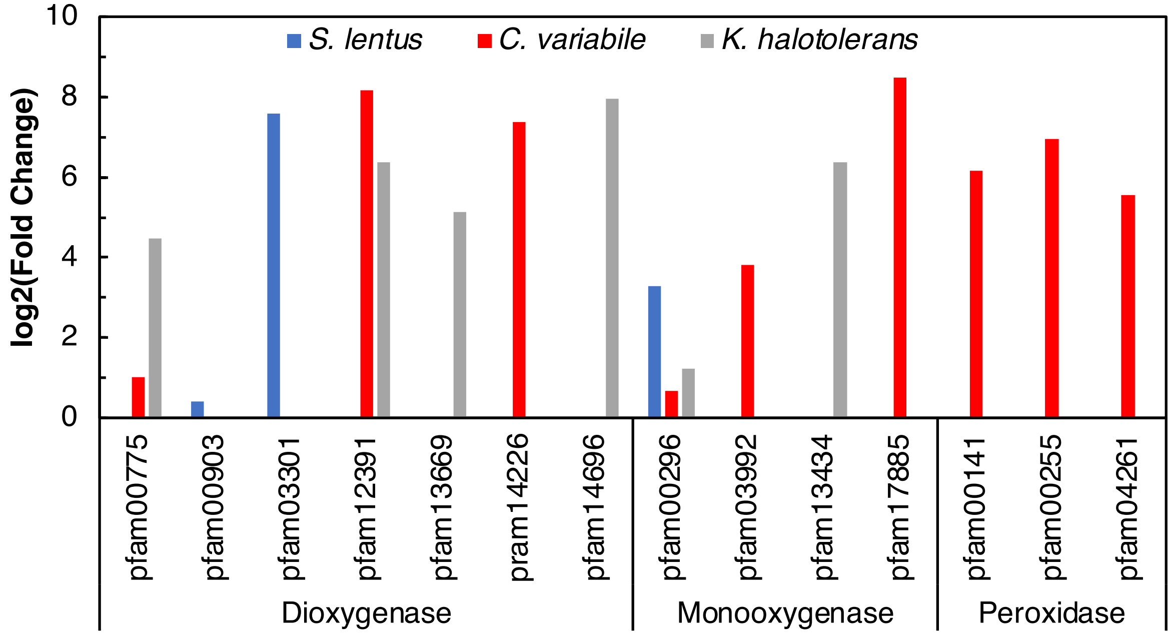


**Supplementary Figure 3**: **Enrichment of alkane metabolism protein families in yellow mealworm gut microbiome isolates.** Families that are statistically enriched relative to the taxonomic Order of an isolate, as measured by a Fisher's exact test, are depicted with their enrichment ($\log_2$(fold change)). All pfams used in this analysis can be found in Supplementary Table 3. Any enzyme in that table that does not appear in this figure was not statistically enriched, $p<0.05$.

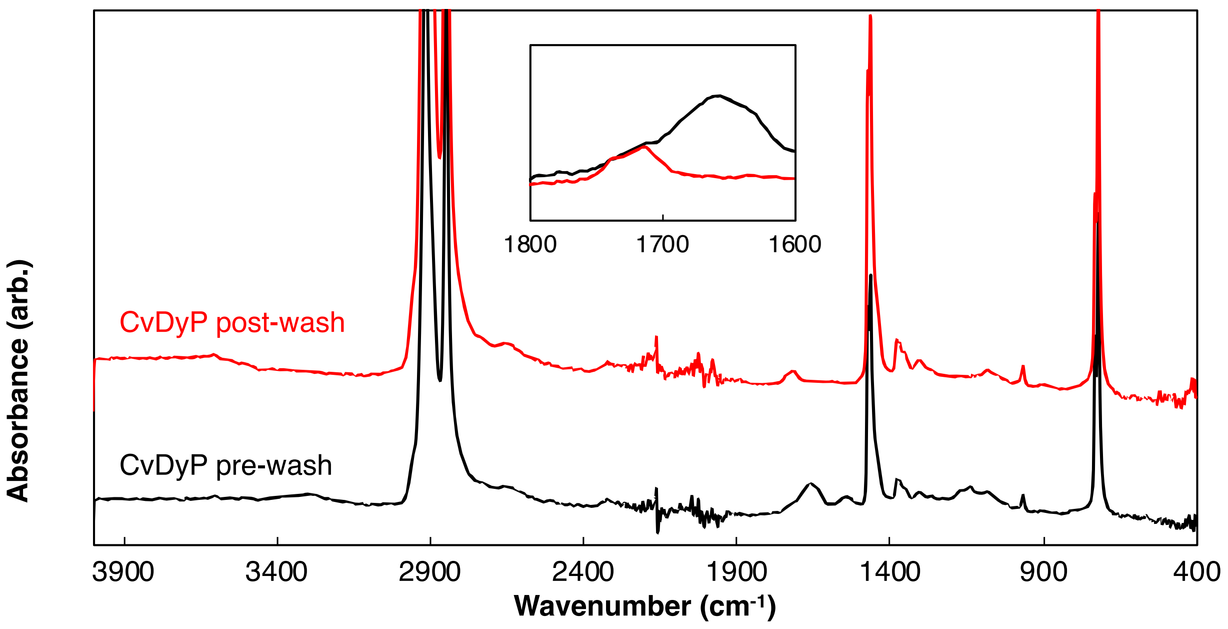


**Supplementary Figure 4: Difference in CvDyP-treated LDPE FTIR spectra pre and post-washing.** FTIR spectra of stripped LDPE treated with pure CvDyP before (black) and after (red) film washing with water and 70% ethanol. Protein contaminant peaks at 3400 $cm^{-1}$, 1650 $cm^{-1}$, and 1550 $cm^{-1}$ are removed as a result of washing and carbonyl peaks between 1700 and 1740 $cm^{-1}$ persist.

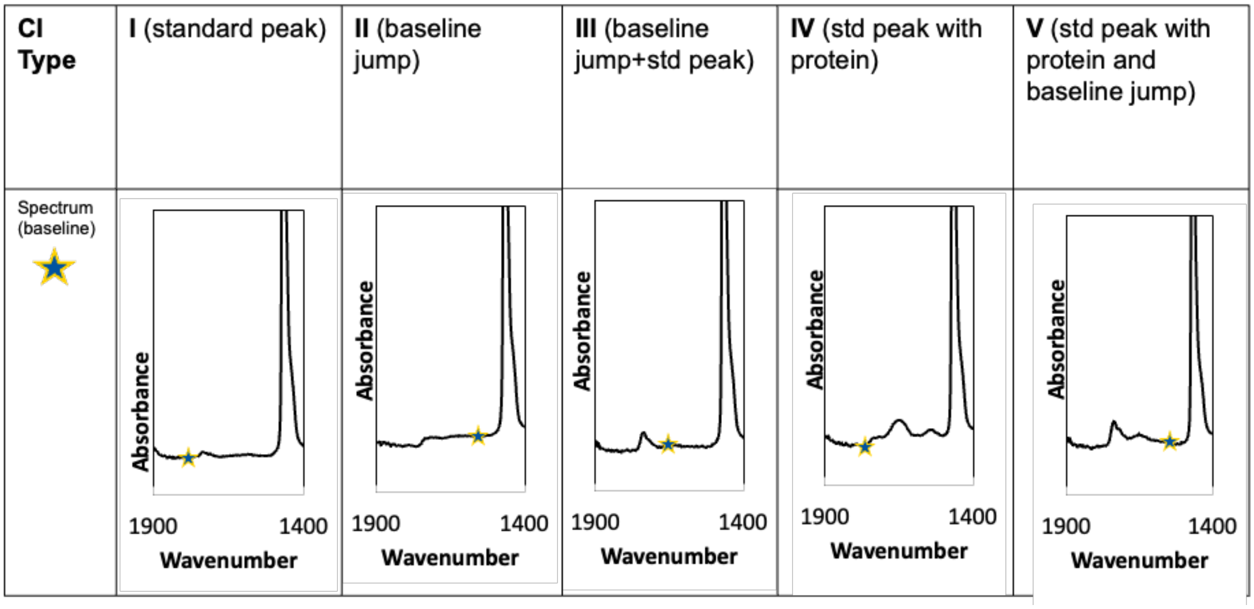


**Supplementary Figure 5: FTIR carbonyl index baseline selection**. When taking the baseline of FTIR spectra, we observed differences that lead to inconsistencies in carbonyl index calculations. In order to use appropriate baselines to calculate the carbonyl index, each spectrum was looked at individually to determine where the true baseline is. There are five types of common biological oxidation spectra that are observed in the carbonyl region (1) standard peak, (2) baseline jump, (3) baseline jump + standard peak (4) standard peak with protein/biomass contamination, and (5) standard peak with protein contamination and a baseline jump. In each case, a different, appropriate baseline value needs taken. This figure gives best practices for defining the appropriate baseline to use when normalizing spectral values.

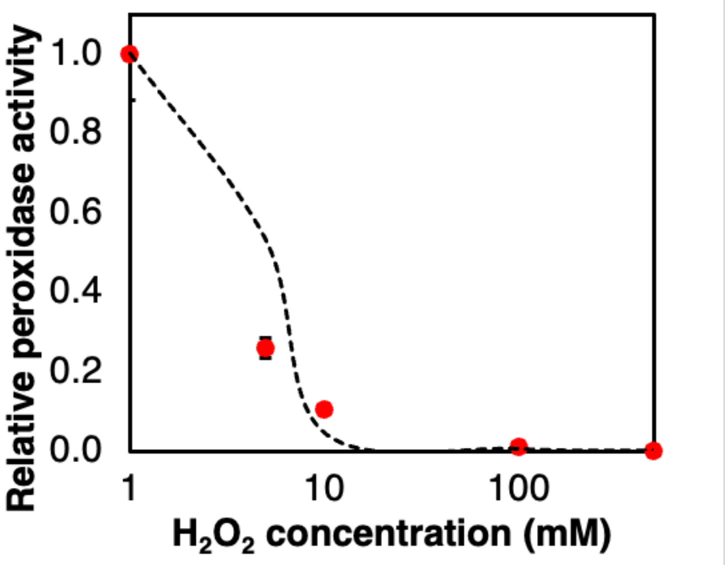


**Supplementary Figure 6: CvDyP activity dependence on $H_2O_2$.** $H_2O_2$ concentration was varied in 300 µL reactions in a 96 well plate at room temperature in pH4 potassium phosphate buffer. Error bars represent the standard error across two biological replicates. The fit curve follows Equation 1 below where $V_{max}$ = 1.0062, $K_I$ = 250, and m = 7.8

**Equation 1:** $$\frac{d[P]}{dt} = V_{max} * \frac{K_I}{K_I + [I]^m}$$

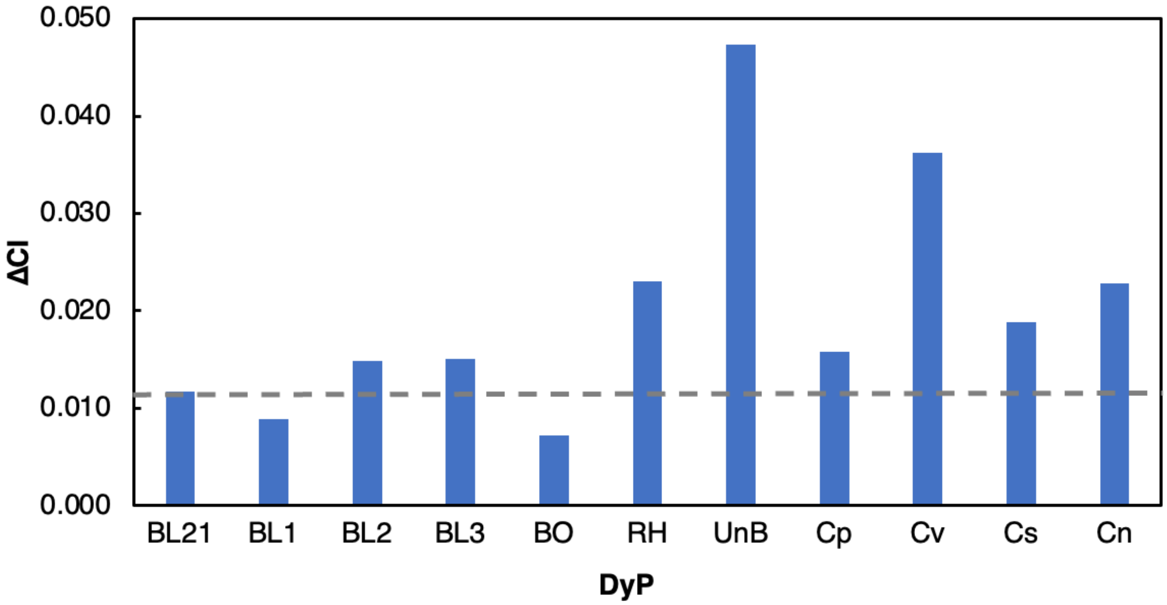


**Supplementary Figure 7: Change in carbonyl index on in-house stripped LDPE films on lysates from *E. coli* BL21 (DE3) over-expression cultures of each tested DyP.** Carbonyl indices were calculated according to Supplementary Figure 4. BL21: E. coli BL21 control, remainder of the bars in the bar chart correspond to the nomenclature codes provided in Supplementary Table 4 for each Type I DyP that was tested.

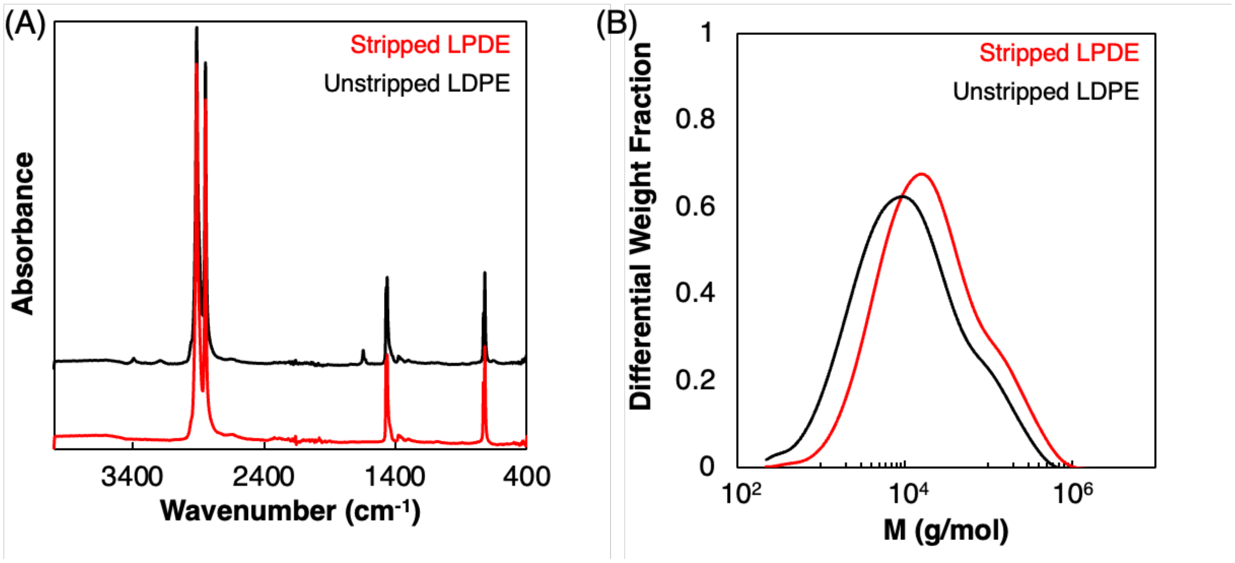


**Supplementary Figure 8: Materials characterization of stripped and un-stripped LDPE.** (A) FTIR spectra showing stripped (red) vs. unstripped (black) LDPE. Unstripped plastic contains additive peaks around 3400, 3200, and 1600 $cm^{-1}$. (B) SEC chromatogram showing molecular weight distribution of stripped (red) vs. unstripped (black) LDPE. In the additive stripping process, LDPE is stripped of lower molecular weight chains, leading to an increase in the average molecular weight.

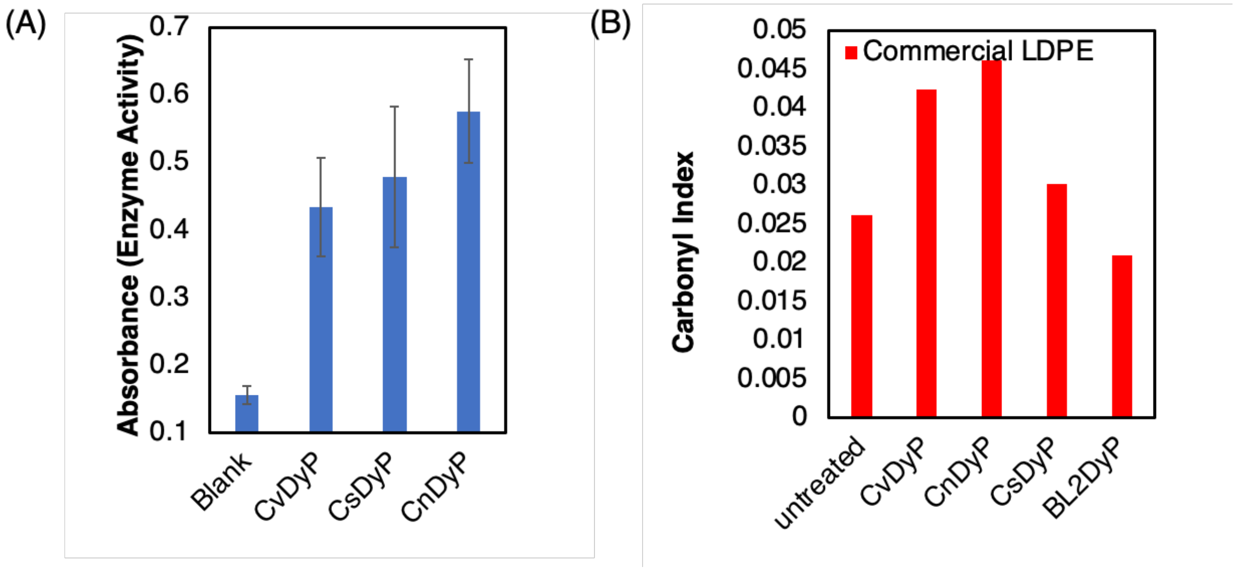


**Supplementary Figure 9: Identified LDPE oxidases are active peroxidases** (A) Pyrogallol peroxidase assay endpoint demonstrating activity of purified class i3 DyPs at room temperature after 24 hours at pH4. Error bars represent the standard error of biological triplicate measurements. (B) Activity of LDPE-active (CvDyP, CnDyP, CsDyP) and LDPE-inactive (Bl2DyP) enzymes on commercial PE films, measured by carbonyl index. CI were calculated according to Supplementary Figure 5.

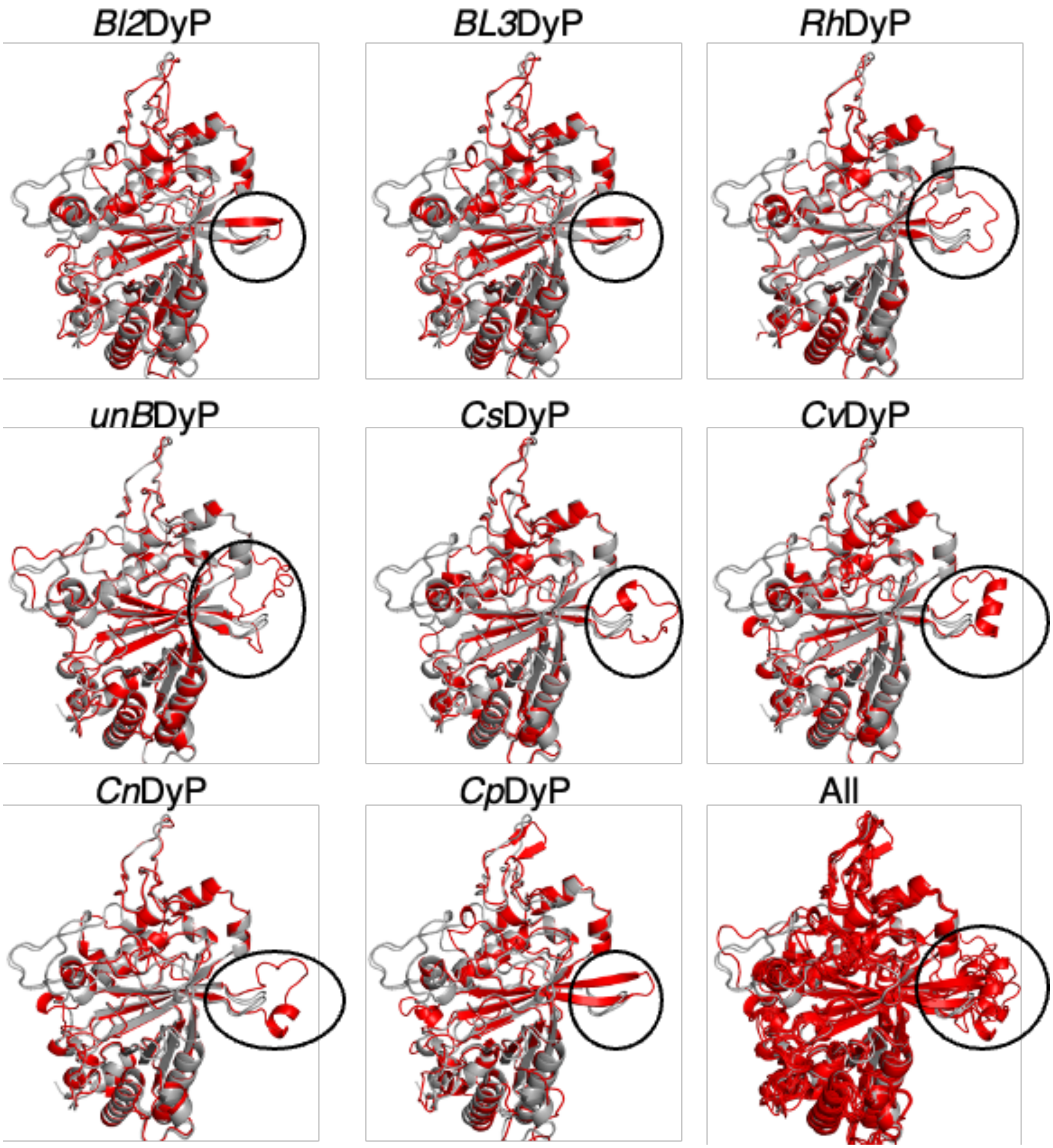


**Supplementary Figure 10: Structural alignment of LDPE-active (red) vs LDPE-inactive (grey) DyPs**. Grey DyPs are inactive BL1DyP and BoDyP in each image. Each label represents the red protein overlaid atop the two inactive proteins in each case. Hydrophobic loop indicated by black circle. Images generated using Pymol version 2.5.2. Enzyme codes are defined in Supplementary Table 4.

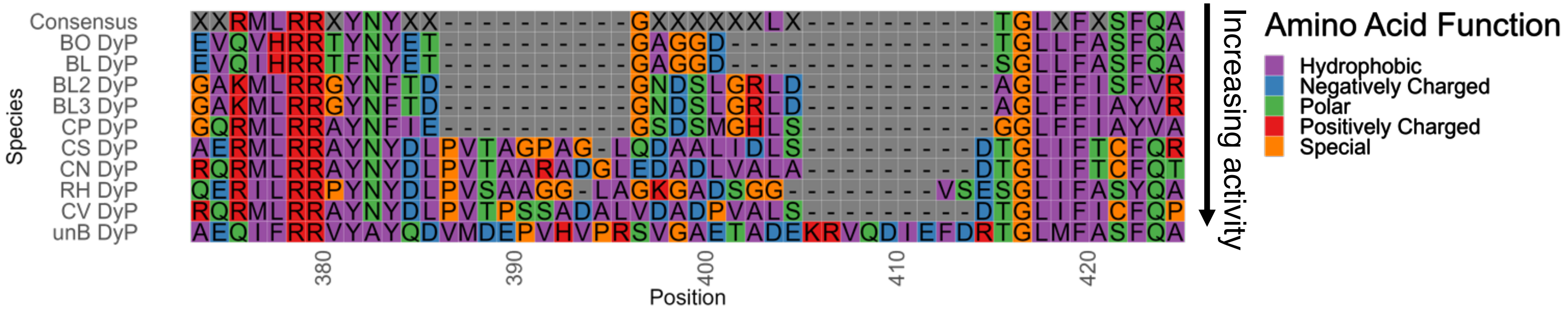


**Supplementary Figure 11: Sequence alignment of hydrophobic loop region across all tested DyPs.** Sequences were aligned using ClustlW. Arrow indicates relative activity level per Supplementary Figure 7. A 50% minimum consensus sequence is provided in the top row.

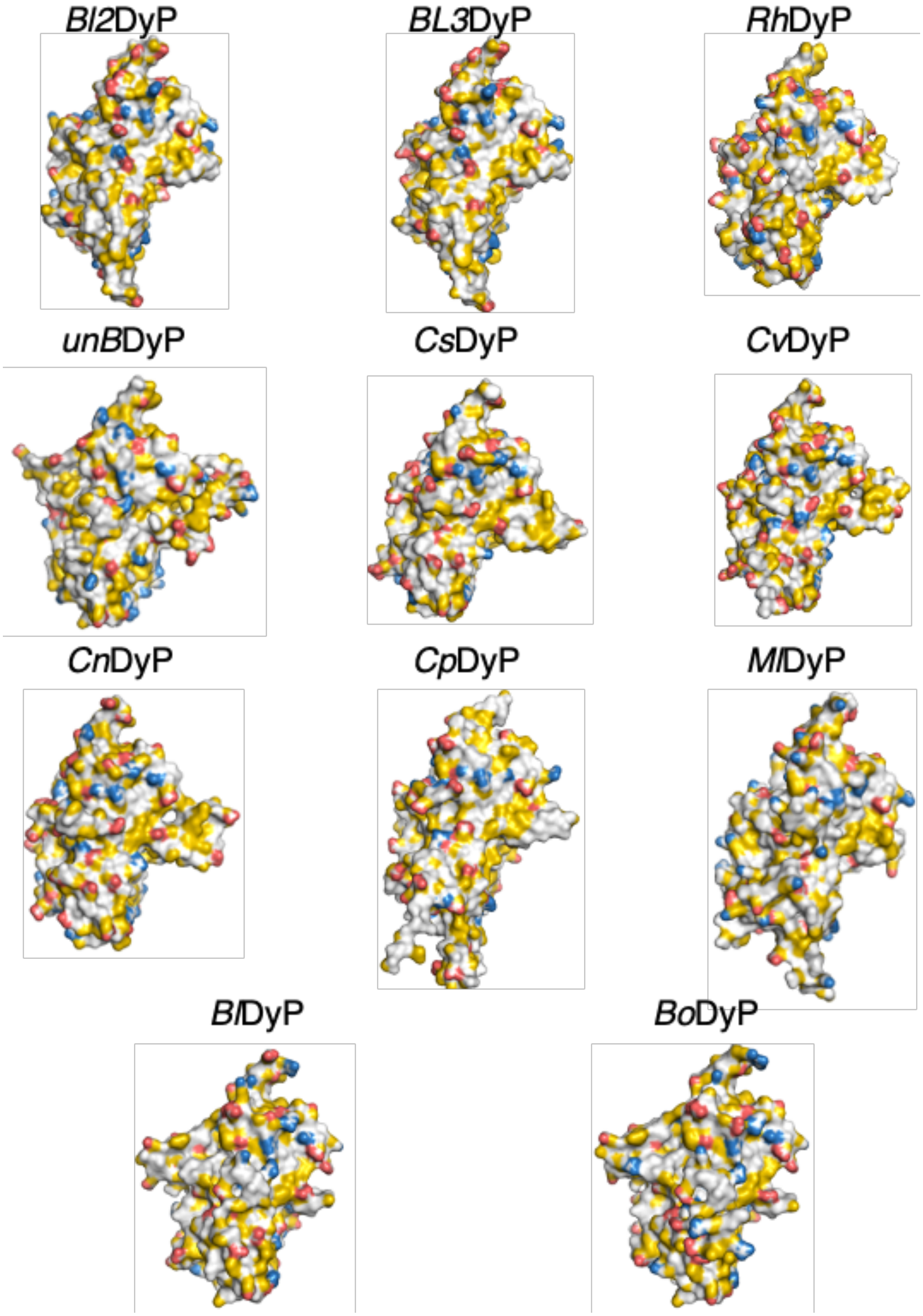


**Supplementary Figure 12: YRB maps of all tested DyPs.** Images generated using Pymol version 2.5.2. Yellow represents hydrophobic atoms, blue are positively charged atoms, and red are negatively charged atoms. Enzyme codes are defined in Supplementary Table 4

(A)

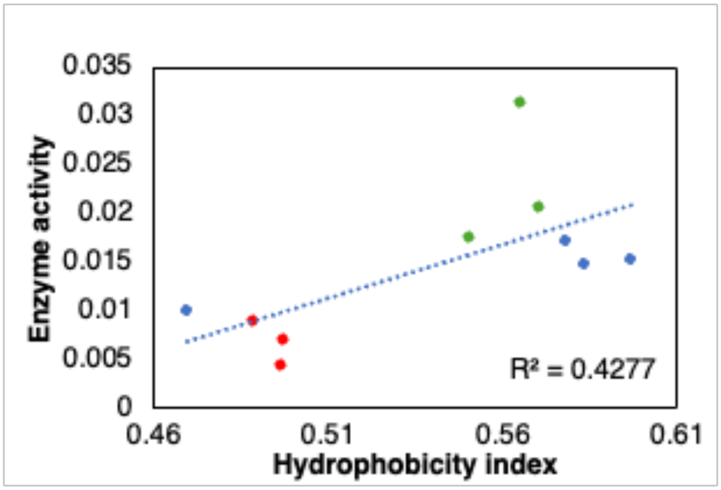


(B)

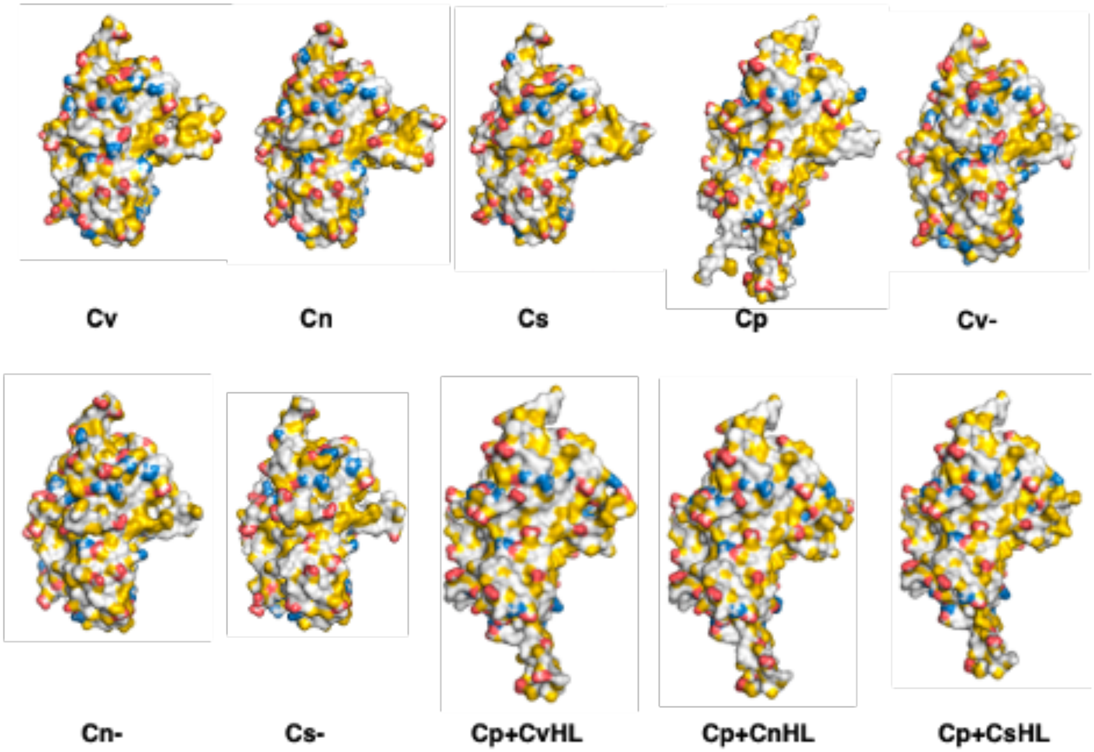


**Supplementary Figure 13: Trends between loop hydrophobicity and LDPE oxidase activity.** (A) Correlation between hydrophobic loop region hydrophobicity and enzyme activity of DyPs in (A). (B) YRB hydrophobicity maps of mutant and native *Corynebacterium* DyPs. Images generated using Pymol version 2.5.2. Yellow represents hydrophobic atoms, blue are positively charged atoms, and red are negatively charged atoms. Enzyme codes are defined in Supplementary Table 4

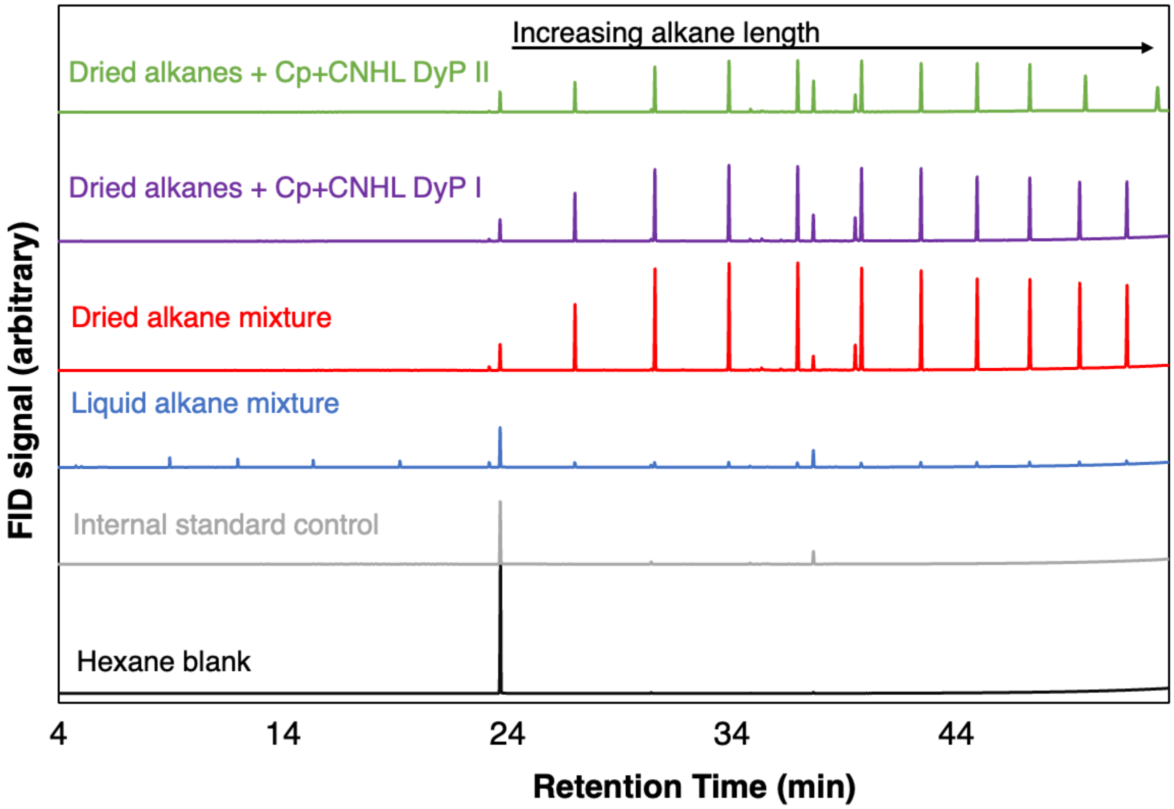


**Supplementary Figure 14: GC FID chromatograms from Cp-CNHL DyP reaction with liquid alkanes.** Control chromatograms were gathered for solvent hexane, internal standard 2-decanone, the vacuum dried C10-C40 alkane mixture, and Cp-CNHL reactions with the dried alkane mixture. Each chromatogram is of a single measurement. Each Cp-CNHL chromatogram represents a single reaction at room temperature, pH 4, after 16 hours of reaction time.

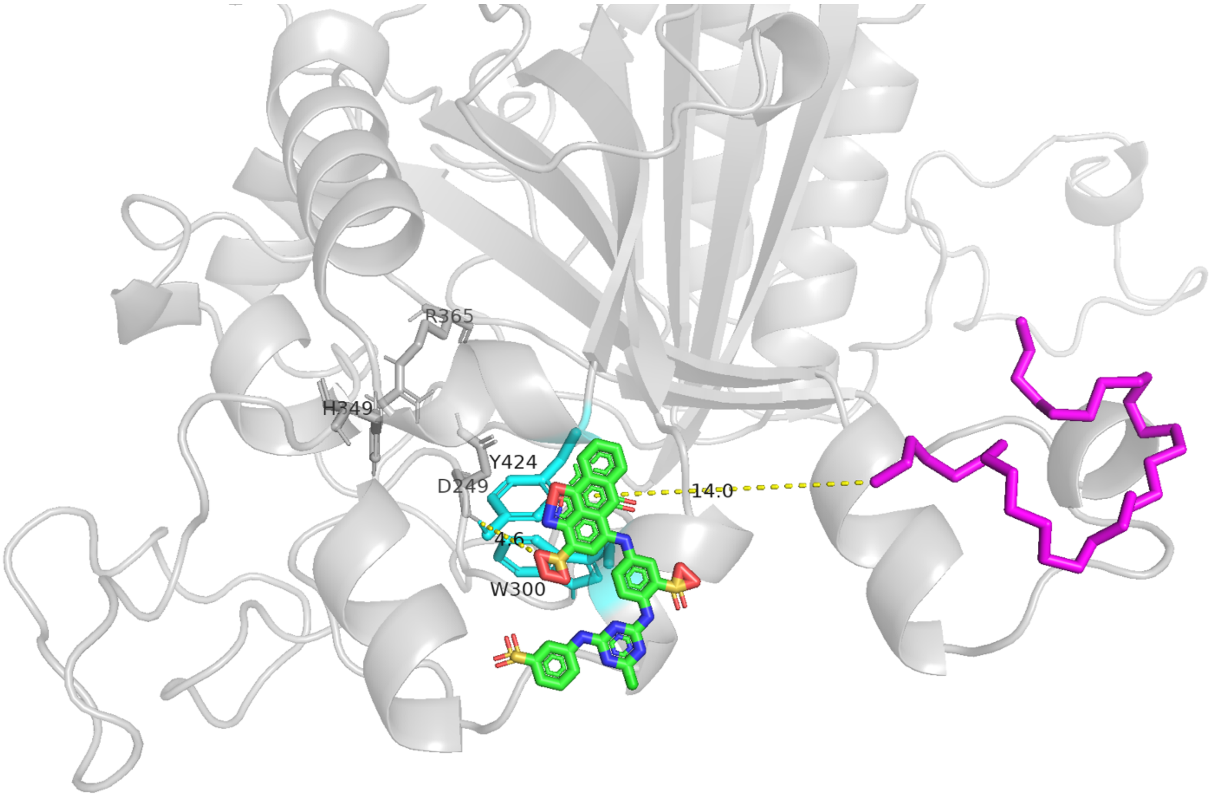


**Supplementary Figure 15: Molecular docking of Reactive Blue 2 to aromatic surface residues and dotriacontane to the hydrophobic loop.** The top pose from Autodock molecular docking simulations of dotriacontane (magenta) with Cp-CNHL and Reactive Blue 2 (RB2, green)) are overlaid. Simulations purposely forced the binding region to the hydrophobic loop region and to key aromatic residues W300 and Y424 (cyan) for dotriacontane and RB2, respectively. This particular binding was done in order to simulate probable locations for proposed binding of each substrate. A non-conservative minimum number of 44 carbons in chain length for LDPE oxidation were calculated by approximating that the dotriacontane molecule was ~14 angstroms from the active site but needed to be 4.5 angstroms away to be oxidized. This remaining distance of ~9.5 angstroms was then used in conjunction with an approximated 5 angstroms per 6 carbons on the dotriacontane chain, estimated by distance measurements in Pymol. Note that this distance represents the absolute minimum number of carbons if the chain is bound in the optimum position on the *Cp-CNHL*DyP loop as shown.

**Supplementary Table 1: List of taxa isolated from mealworm guts under various feed conditions**

| ISOLATE NUMBER | TAXONOMIC IDENTIFICATION | MEALWORM FEED | ISOLATION MEDIUM |
|---|---|---|---|
| **1** | *Staphylococcus lentus* | Polypropylene | YPD |
| **2** | *Enterococcus termitis* | PS+oats+antibiotic | LB |
| **3** | *Staphylococcus galinarum* | PS+oats+antibiotic | LB |
| **4** | *Listeria fleischmanii* | PS+oats+antibiotic | TSA |
| **5** | *Enterococcus sp.* | Ps+antibiotic+antifungal | Pseudomonas |
| **6** | *Mangrovibacter yixingensis* | Ps+antibiotic+antifungal | Mackonkey |
| **7** | *Corynebacterium variabile* | HDPE+antifungal | Pseudomonas |
| **8** | *Enterococcus faecalis* | PP+antibiotic+antifungal | YPD |
| **9** | *Brachybacterium nesterenkovii* | PP+antifungal | Blood |
| **10** | *Brevibacterium epidermidis* | PP+antifungal | TSA |
| **11** | *Lactococcus petuari* | HDPE+antifungal | Blood |
| **12** | *Lactococcus garvieae* | PS+antifungal | TSA |
| **13** | *Kocuria halotolerans* | LDPE+antibiotic+antifungal | LB |
| **14** | *Klebsiella aerogenes* | HDPE+antibiotic | LB |
| **15** | *Intestinirhabdus sp.* | HDPE | Medium B |
| **16** | *Enterococcus gallinarun* | HDPE+oats | Blood |
| **17** | *Escherichia fergusonii* | PS+oats+antibiotic | Blood |
| **18** | *Enterococcus casseliflavus* | HDPE+oats | Blood |
| **19** | *Enterococcus thailandicus* | LDPE+Oats | TSA |
| **20** | *Aspergillus fumigatus* | HDPE | Medium B |
| **21** | *Enterococcus sagiogensis* | PP | Beef medium |

**Supplementary Table 2: Location of maximum carbonyl peaks by each isolate on LDPE films**

| | *Carbonyl peak location ($cm^{-1}$)* | *C=O functionality* |
|---|---|---|
| *S. lentus* | 1745 | Ester* |
| *E. termitis* | 1735 | Aldehyde |
| *C. variabile* | 1712 | Ketone˙ |
| *B. epidermidis* | 1735 | Aldehyde |
| *L. garvieae* | 1737 | Aldehyde |
| *K. halotolerans* | 1729 | Aldehyde |

**S. lentus* spectrum has a secondary peak at 1712 $cm^{-1}$ indicating a ketone*

*˙C. variabile spectrum has a secondary peak at 1735 $cm^{-1}$ indicating an aldehyde*

**Supplementary Table 3: putative alkane metabolism pfams from comparative genomic analyses that yielded positive hits from JGI IMG. These enzyme families were present in genomes from the Order of each of the five tested isolates, *Staphylococcus lentus, Enterococcus termitis, Corynebacterium Variabile, Brevibacterium Epidermidis,* and *Kocuria halotolerans***

| pfam | Function |
|---|---|
| **pfam00141** | Peroxidase |
| **pfam00255** | Glutathione peroxidase |
| **pfam00296** | Luciferase-like monooxygenase |
| **pfam00351** | Biopterin-dependent aromatic amino acid hydroxylase |
| **pfam00743** | Flavin-binding monooxygenase-like |
| **pfam00775** | Dioxygenase |
| **pfam00903** | Glyoxalase/Bleomycin resistance protein/Dioxygenase superfamily |
| **pfam01083** | Cutinase |
| **pfam01231** | Indoleamine 2,3-dioxygenase |
| **pfam02332** | Methane/Phenol/Toluene Hydroxylase |
| **pfam02578** | Multi-copper polyphenol oxidoreductase laccase |
| **pfam02668** | Taurine catabolism dioxygenase TauD, TfdA family |
| **pfam02900** | Catalytic LigB subunit of aromatic ring-opening dioxygenase |
| **pfam03060** | Nitronate monooxygenase |
| **pfam03098** | Animal haem peroxidase |
| **pfam03150** | Di-haem cytochrome c peroxidase |
| **pfam03241** | 4-hydroxyphenylacetate 3-hydroxylase C terminal |
| **pfam03301** | Tryptophan 2,3-dioxygenase |
| **pfam03992** | Antibiotic biosynthesis monooxygenase |
| **pfam04116** | Fatty acid hydroxylase superfamily |
| **pfam04209** | homogentisate 1,2-dioxygenase |
| **pfam04261** | Dyp-type peroxidase family |
| **pfam04444** | Catechol dioxygenase N terminus |
| **pfam04454** | Encapsulating protein for peroxidase |
| **pfam04663** | Phenol hydroxylase conserved region |
| **pfam04989** | Cephalosporin hydroxylase |
| **pfam05118** | Aspartyl/Asparaginyl beta-hydroxylase |
| **pfam05373** | L-proline 3-hydroxylase, C-terminal |
| **pfam05721** | Phytanoyl-CoA dioxygenase (PhyH) |
| **pfam05995** | Cysteine dioxygenase type I |
| **pfam06052** | 3-hydroxyanthranilic acid dioxygenase |
| **pfam06234** | Toluene-4-monooxygenase system protein B (TmoB) |
| **pfam06537** | Di-haem oxidoreductase, putative peroxidase |
| **pfam07746** | Aromatic-ring-opening dioxygenase LigAB, LigA subunit |
| **pfam07976** | Phenol hydroxylase, C-terminal dimerisation domain |

| | |
|---|---|
| **pfam10014** | 2OG-Fe dioxygenase |
| **pfam11723** | Homotrimeric ring hydroxylase |
| **pfam11794** | 4-hydroxyphenylacetate 3-hydroxylase N terminal |
| **pfam12391** | Protocatechuate 3,4-dioxygenase beta subunit N terminal |
| **pfam13434** | L-lysine 6-monooxygenase (NADPH-requiring) |
| **pfam13669** | Glyoxalase/Bleomycin resistance protein/Dioxygenase superfamily |
| **pfam13794** | tRNA-(MS[2]IO[6]A)-hydroxylase (MiaE)-like |
| **pfam14226** | non-haem dioxygenase in morphine synthesis N-terminal |
| **pfam14696** | Hydroxyphenylpyruvate dioxygenase, HPPD, N-terminal |
| **pfam15461** | Beta-carotene 15,15'-dioxygenase |
| **pfam17885** | Styrene monooxygenase A putative substrate binding domain |
| **pfam18331** | PKHD-type hydroxylase C-terminal domain |
| **pfam19298** | 3-Ketosteroid 9alpha-hydroxylase C-terminal domain |

**Supplementary Table 4: Enzyme and sequence information from screened *Corynebacterium variabile* enzymes:**

| ID (IMG ID) | Pfam | Cofactor(s) | Sequence |
|---|---|---|---|
| P1 (Ga0530663_0133_ 26266_26778) | 00225 | $H_2O_2$ | MTENTPDTTDIKSIPATTLDGDPLDLHDFDGPLLIVNVASKCGLTPQYTALEQL QKTYSTTDGTGLTFLGVPCNQFAGQEPGSAAEIQTFCSTTYGVTFPLLAKTD VNGDDRDPLYQALTTTQDAEGEAGDIQWNFEKFLVSADGRVLNRFRPQTTP DDPAVIDAIEAAL |
| M1 (Ga0530663_0122_ 22947_24377) | 00296 | FMN, $Mg^{2+}$, NADH | MTRIRFNAFDMNCVVHQSPGLWRHPEDHSREYTDLRYWTDLAQLLETGYFD GLFLADVLGLYDLYGASPDAALRAATQTPVNDPLPLVSAMAAVTDNLGFGLT TGTAFEHPFPFARRISTLDHLTDGRLGWNVVTGYLPSAAENTQSRGQLDTFE HDARYDHADEYLEVLYKLWEGSWEDDAVLADPVSGVYTDPTKVRPVNHTGT HYTVPGIHVSEPSPQRTPVIFQAGASSRGSTFAATHAEAVFVAAPTQEALAST VRTLTDKLADAGRSREDVVIYALLTVITDATEDAARAKAAEYRSYIDHEGALAL MSGWMGVDFSTFDLDDPVAGIESNAIQSAVAAFQRSAGEEGREWTVRELAE FGGIGGLGPVVVGSGESVADQLEEWVEATGIDGFNLAYAVTPGTFRDVIDHV VPVLQARGEYPTAYAPGTLREKLFADRTGAGPHVPHNHSAARYRPALTASPT SVPTGAHP |
| P2 (Ga0530663_0293_ 23766_25013) | 04261 | $H_2O_2$ | MDTNFRVTRRGFLAGLGLVSATAAVGATTACAGDSDPDPVSNTDTTVPFDG PHQAGIATPLQRHNTTVAFTLRDGVDATGIRRLLRIWTGDARRLTQGEPVLAD LEPELASDPGRLTVTCGFGRGLFEKARLTDKSPDWLEPLPGFAGDRLEARW GDRDLVLQICGEDRTTVSHALRVLVRGGADYARPSWTQTGFLDIPQDADGT PGTPRNLFGFKDGTVNPRTDAEFDDQVWISGEAATHPSQIGGTCMIIRRIAFD MPLWESADRPTREVSMGRTIVEGNPLSGEKEHDDPDTAAVGADGLPVIDRN AHIALAGVHDGDTRQRMLRRAYNYDLPVTPSSADALVDADPVALSDTGLIFIC FQPDPRTSFIPVQRRLAAGDRLNEWITHVGSAVFLVPAGTTEEEYWGEALLG |
| M2 (Ga0530663_0069_ 83329_84399) | 00296 | NADH | VKAFGFLSFGHYDLSTSGRMPGAKENLKNALEIAVGADELGVNGAYFRVHHF APQHSAPMPLLSAIAARTSNIEVGTGVIDIRYENPLQLAEEAAQLDLLSDGRVA LGVSRGSPEPVDRGYEAFGYHPAADDPKGAAMAREKFERFMAAVDGEGMA TSLPLEEQYPRMCHPNVPVPVMPHSMSLRKHVWWGAGSRDTAVRAAHDG VNLMSSTLLTEANGDRFEDLQAEQIRAYRAAWKEAGHQWTPRVSVSRSIIPV TSEEDMRMFGIGGDEGGIGYIDDGQATFGRTYTDEPDKLVEQLKKDAALTEA DTLMITIPSQLGVDANLRILQSFAEHVAPELGWEPNTAGTVTGYPLV |

| | | | |
|---|---|---|---|
| M3 (Ga0530663_0122_ 24374_25753) | 17885 | $FADH_2$ | MSTQHTPDNLSVENAQTTPAGAPRSIAVIGAGLAGAGAALVLARAGYRVDLY<br>SDRTREALRDEVPATGTAILFGASRTADAKIIDDLYAGSPAAHFGASAAGLAN<br>PDNPSGVDSFPARYTYEAQSVDARLRADDRLAAFLDLTGDTASAFHVTRVTP<br>ADLDRIAAEHDLTLVATGKGGLADAFTTDHERTPYPGPQRRVLTVTAAGLSS<br>DHLFSGRAEVPGVNNALLSLHGEGEVFVGPYLHKDGIDAWVILAWAKPGTAT<br>EKAFAEATDAESSLRILQDLHHRVLPDVADDLDRLRPIDADPHSWLAGAVTPR<br>VRRPVGYTDGGNLVAALGDTAIAVDPIAGQGAQLGDFQVAALVEGLTAAEAS<br>GQPWDADLLTDLFETHWAGHGAAGVAVTSLFLGDPLYAEVAGAFFGNAGTD<br>PAYATALFSLFSEPAPALTLRTAEDVAALAASFRNSTAKVTA |
| D1 (Ga0530663_0126_ 51261_52328) | 14226 | $Fe^{2+}$ | MVQTTTPTDNATVTDTNAELTLESTMGGIGTETTDRSVPVIDLDNWDLTDEQ<br>WDAKAEEFWDAATTIGFFQLKNFGITRAEIEDAFATSARFFALPKETLETVAKP<br>KGRNVGFEHKSQIRPSTGTPDEKESYQITRPLMDGLWLDEDANPSIDGFKDA<br>SLAFEAKCHEVAMRVLEFFAVKLGFERDYFRKVHNPASDLHQCTLRMIHYMA<br>LDAKDNVADPNGNPVWRAGAHTDFNLLTLLFQTDGQSGLQVMPGADAGQD<br>VQAWTPVPAFTDVLTCNIGDMLMRWSDDRLKSNFHRVKAADIGVDIPERYS<br>MPYFAQADRDAVIAGPEGRYEPMTAGEYLDMRIQANFGKAESTK |
| D2 (Ga0530663_0230_ 79218_79910) | 12391 | $Fe^{3+}$ | MIPISNATVDGSFAPLHFPEYRTTVLRNPSNDLLMVPQRLGELSGPVFGAADL<br>RGEDNDLTKVNGGEAIGQRIFVHGRVTGEDGRPVPETLIEVWQANSAGRYR<br>HKNDSWPAPLDPHFNGVGRTLTDKDGNYSFYTVQPGCYPWGNHHNAWRP<br>AHIHFSLFGRQFTERLVTQMYFPGDPMFFQDPIYNSVPAGARERMISVFDYD<br>ETRENYAMGFRFDIALRGRNATPFE |
| P3 (Ga0530663_0083_ 164432_166714) | 00141 | $H_2O_2$ | MTDPMSGCPVAHGGGHTPQGGHAPQESQAERGEAQGSRLPLPTEGNANQ<br>RWWPKRLNVRLLAHNPQEVRPTPADFNYAEAFSQLDLPQVKKDIEDVLTTSQ<br>SWWPADFGHYGPLIIRMAWHSAGTYRSHDGRGGGGAGQQRFAPLNSWPD<br>NVGLDKARRVLWPVKKKYGQNLSWGDLMILAGNVALESMGFETFGYSGGR<br>EDVWEPEDVYWGSETDWLGTDARYSGTNDTSRDLQKPFGATTMGLIYVNP<br>EGPEGNPDPAAAAHDIRETFGRMGMNDEETVALIAGGHTFGKTHGAAPDNE<br>ENVGPDPEEASLDRQGLGWQNHHGEGKGDDQITSGLEVTWTYHPTRWDN<br>EFFHILFAYEWELTTGEGGHFHWRPKDGGGADMVPMAHSAGRREPRMLTT<br>DISLRVDPSYEKISRRFKDDQNALNDAFARAWFKLTHRDMGPKSRYLGPEVP<br>AEDLIWQDPLPDAEGDPVTPQDVEALKKTVLDSGLSVSQLVKTAWASASSFR<br>GTDFRGGANGARIRLEPQRSWEANEPARLAEVLTALEAIRTEFNENNAPRHL<br>SLADLIILAGNAAIEKAAADAGTDVSVPFTPGRVDATQEQTDVDGFSYLEPQA<br>DGFRNYQNADLDIPAEHLLIDKADLLGLTAPQMTVLVGGLRVLGATYGSTGH<br>GVLTDRPGVLSTDFFVNLLELGNDWVATDDNATVFRCTDGETGEEKWTGTR<br>ADLVFASNSELRAQAEVYASDDAAEKFIADFVAAWTKVTEADRFDLHG |

| | | | |
|---|---|---|---|
| D3 (Ga0530663_0127_2496_3656) | 00775 | $Fe^{3+}$ | MGDVPGIRSGKTECHPIHTGLTAGLTVREPRPQGVTSMSTSNSPEPTQTPMIDSDPNGPFRYPVSDIRDQDEATPGIMPSQTVGPYVHIGLTWPGAENMVEEGTDGAVEVTFQVTDGAGHLIKDAMIEIWQAGSDGVYPSPLDPRAGEDAGREGFRGLGRGMCDSETGLVTFRTVRPGAVPAPDGGTEAPHLKVGVFARGMLERLYTRLYFPEQSEANDADPVLNAVPEERRDLLVASTVDGDADSYRMDIVMQHEDATRETPFFALEGPAFEGRLLDRPDEDVVDQGPAFDIRTLMTRRTVLGVGAAGVGAVALAACSDNSSSSAATTSASSSASSSASVTDLPDAEIPDETAGPYPADGSNGPDVLEESGIVRADIRSSIGADDPVDGVPLEFSITVTDMANDDVPFENVAVYAWHCNAEGLYSMYSEGVEDETWLRGVQVADADGTVSFTSIVPACYTGRWPHIHFEVYPSVDDITDSENAIATSQIAIPEDVCNTVFALDAYEGSSENLSQVTLESDNVFGDDSGVSQLATMSGDVDSGYIASLTARVDTTTEPTGGAAPEGGPGEGGPGGEGGPGGEGGEPPAGEPPAGSPGAIASTN |
| M4 (Ga0530663_0083_53148_53471) | 03992 | NADH | MILINVRFRPLPEYVENFREHVADYTAACRAEEGCLFFEWYRNTDDSDEYLLVEAYKDGADVAHVQSDHFKASCELFPTLLSETPQIINDHIDGKTEWDRMAEFSVD |

**Supplementary Table 5: Sequence information of class I DyPs tested for LDPE-oxidase activity:**

| Organism | Code | Sequence |
|---|---|---|
| *Brevibacterium linens* | BL | MVTTEPSRTGFSRRGLLTSAAAAGGAGLVGATAGFALGRGPGSQGTGTGDGGLPRRDLDGANGPQ<br>TEPFFGTHQSGVETPTQAYAHFIAFALKPGIKAAEAVRWLRLLTADAAALTQGQAPLADSESELAVDP<br>ARLTVTFGFGGKLVALAGKEHVPDWLKPLEKFSIDRLDADRSKGDLLLQICGDDPLTLAHARRMLFKD<br>SRSFAEVAWQRDGFRRAYGSSGEGKTQRNLFGQLDGTANPGPGSEDFARIIWGQGTETTDPVFSA<br>RGEPPADLGAHLPAWMRGGTTLVLRDIAMNLDTWDKADRPAREFSVGRTMDTGAPLSGKDEFDVP<br>DFTAVDERGLTKISQVAHIARARDGLGPEVQIHRRTFNYETGAGGDSGLLFASFQADIERQFLPIQRRL<br>AEVDLLNEWTTPIGSTVWAIPPGATEDGYVGQELFEG |
| *Brevibacterium linens* | BL2 | MSTDEHSSSADGAVRGIPRRRLFALGGAAGALGIGGLGGGAVGYAVRGAQEKKQSAVHLSYPFRGE<br>KQQGILTPAQDNMYTAAFDVTTTDRQALISLLEDWTVAAEQMSAGNLVGGAPDANAELPPKDSGEV<br>WGYPASGLTITFGFGATLFVDTDGKDRFGLKDKMPAILKEGVPKFANEALHADASNGDLLVQACAND<br>PQVAVHAIRNLTRIAFGTAKLRWGQIGYGRTSSTSTQQETPRNLFGFKDGTNNIKSEDPQKELEKHLW<br>VQSGDDPASDSWLAGGTYFLARKIHMYLEIWDRVSLAEQEDIIGRDKRFGAPQSVAEPDTDEEFTPL<br>DFSAQNAEGAPAIDARAHVAIVAPEHNKGAKMLRRGYNFTDGNDSLGRLDAGLFFISFVRDPRTNFY<br>PILKTMAQKDLLTEYLQHRASALFAIPPGVGTGDTMIGQKLFS |
| *Corynebacterium provencense* | CP | MPRGSVSRRGFLTGLGLTGLGAAAAGTAGAASLASCSSGEDDRNAGQTVPFRGDHQAGVTTAQQE<br>HLHIVAFNVLTDDRDELRDMLATWTTMAERMTRGEQTTDGGALTGTDADANTDPDGTLNLVPEDTG<br>EAVDLSAGRLTVTVGYGPSLFDGRFGLQDRRPAELEPLPKFPGDQLVPDLCDGDIVIQACSDDPQVA<br>VHAVRNLTRAGSGVVEVRWSQLGYGAASRTTKEGDTPRNLFGFKDGTRNITADETDDINNMVWVSA<br>DRPDSAGPGGWMTGGTYMCVRRIRMLLEVWDRQILDDQQHTFARYKGSGAPIGAHDEFDDLPFDLY<br>LGTGGPAIPRDSHVYLAHPDQNNGQRMLRRAYNFIEGSDSMGHLSGGLFFIAYVAAPSVNFTPVQMK<br>LAHEDKMNEYVRYESKAVFACPPGLAEGSSANWGTALFGG |
| *Rothia halotolerans* | RH | MTADRKGPGSGRKNLLTRRNALIGGATATAGAAAAVGIDTVRRGGLPGGGGNPQVEDASSTAPLAE<br>ARVGFHGERQAGVVTPAPAYGNFIAVDLREDVGRADLRRMLKVLSADAADLAEGLPPIADQEPELTEI<br>PANLTVTVGFGERVFDVVDPSAKPGWLKPLPAFEQIDRLREEYSDGDLLLQICSDDRMTLAHAQRLLL<br>KGLRGFGTVRWVQEGFRNASGSLKEGTTMRNLFGQVDGTINPTTEDSSMDDVVYGLMDGLEPWAP<br>GGTSLVIRRIHMNLDTWDEADAPGREDAVGRKLSNGAPLTGEREEDPADLSATTPLGFHVIADYAHIR<br>RATATTPQERILRRPYNYDLPVSAAGGLAGKGADSGGVSESGLIFASYQADPVKQFLPIQRRLAELDM<br>LNMWTVPIGSCVFAIPPGCSPGGFVGDFLFEG |

| | | |
|---|---|---|
| *Brevibacterium linens* | BL3 | MSNDEPIEAGSTEGKVRGIPRRRLFALGGATGALGIVGVGGGAVGYAVRGAQEKKQSAVQLSYPFRA<br>EKQQGILTPAQDNMYTVAFDVTTSDRQALIGLLEDWTVAAEQMSAGDLVGGAPDANAELPPKDSGE<br>VWGYPASGLTITFGFGATLFVDEAGKDRFGLKGKMPQVLKDGVPKFANEAIHASESNGDLLVQACAN<br>DPQVAVHAIRNLTRIAFGTAKLKWGQIGYGRTSSTSTEQETPRNLFGFKDGTNNIKSEDADGELDKHL<br>WVQSGDDPASDGWLTGGTYFLARKIHMYLEIWDRVSLAEQEDIIGRDKRYGAPQSVAEPTADEEFTP<br>IDFSAKNAEGAPAIDARSHIAIVAPEHHKGAKMLRRGYNFTDGNDSLGRLDAGLFFIAYVRDPRTNFYP<br>ILKTMAQKDLLTEYLQHQASALFAVPPGIGTGDTMIGQKLFT |
| *Brevibacterium ocianii* | BO | MTTEPERSGFSRRGLITSAAAAGGAGLVGATAGFGIGRYGPGSNGASDGEGGQPRRDLDGANGAE<br>VEPFYGTHQSGVETPPQAHAHFLAFTMKPGIKAAEAVRWLRLLTADAAALTQGRPPLADSESELTVD<br>PARLTVTFGFGPGLVALAGKQHVPDWLKPLEKFRIDDLDPDRCTGDLLLQICGDDPLTVAHARRMLFK<br>DTRTFAEVAWQRGGFRRAYGSTGEGKTQRNLFGQLDGTANPGPGSEDFARIIWGQGTDRADPVYS<br>ARGEPPTDLGSHQPPWLDGGTTLVLRDIAMNLDTWDKADRPAREFSTGRTLDTGAPLSGTDEFDVP<br>DFTAVDKRGLTKISKAAHIARARDGFGPEVQVHRRTYNYETGAGGDTGLLFASFQADIERQFLPIQRR<br>LAEVDLLNEWTTPIGSTVWAIPPGTSDGGYIGQELFEG |
| *Unclassified Brevibacterium* | unB | MTTDAPNPSGVPDTAGVTRAADVTDAADQRESRGPNRRRVLTGTAAGAAAGLVVGAGGGVAGGRA<br>WGSATAGGPAGATLDTAGASGGQTVAFYGRHQPGVETPPQAHARFVALDLRRGVGAADAVRMLRL<br>LSDDAAALMAGRAPLADSEPELAGRPARLSITFGFGPGLVSLAGPRARPAWLAPLPAFPVDRLQKSL<br>SRGDLLLQVCCEDPLTLSHAVRMLLKDARSFARIAWTQAASRRAYGTDPAGTTIRNPFGQVDGTANP<br>VPGTPAFDRLVWGESSGGAEGVDWGPRGQPPADLGAGLPKWLHGGTTLVLRDIAMNMATWDEAD<br>GPAREFSVGRRLSNGAPLTGREEHDVPDLGATDSVGFTVISPSSHVARSRNESDPAEQIFRRVYAYQ<br>DVMDEPVHVPRSVGAETADEKRVQDIEFDRTGLMFASFQADVTRQFLPIQRRLGEVDLLNQWTTPIG<br>STVWAIPPGAAGAGDYVGSTLFG |
| *Corynebacterium neomassiliense* | CN | MKRHSSSGQLSRRSFLTTVGLVSATAAAGVTTACASGEAGTDGSADQQGADRPDPVVPFDGPHQA<br>GIATPVQAHNTTVAFTLRDGVGPKEVRRLLNIWTSDARRMCSGEPVLADLEPELSSDPGRLTVTCGF<br>GRGLFTAAGVADRAPRWLGPLPAFSTDRLDPRWGERDLVIQVCGDDRTTVSHALRILVRGGADYAR<br>PSWTQTGFLDVPVGPDGEPGTPRNLFGFKDGTVNPRTETDFDEQVWIDDAEASHPAHVGGTCMVIR<br>RVAFDMPLWESADRTTREVSMGRTITEGAPLTGRAEFDEPDLAAIGDDGLPVIDAHAHIALAGVHDGD<br>TRQRMLRRAYNYDLPVTAARADGLEDADLVALADTGLIFTCFQTDPRTSFIPVQRRLADGDRLNEWIS<br>HVGSAVFFIPPGTAEGEYWGEKLLG |
| *Corynebacterium sp.* | CS | MPGTSPHRVTRRGFLGSLGTLGVLGASAAVTTSCAQDAGGTQDAAASGAAGPSSVPFDGAHQAGV<br>ATAPQAHNTTVAFTLRDGADRTTVQRLLRIWTGDARRLCSGDPVLADLEPELASDPGNLTVTCGFGR<br>GLYEAAGIADKAPSWLKPLPHFTGDALDDTWGGRDVVLQICGDDRTTVSHALRVLVRGGADHARPS<br>WSQTGFLDIPVHDGDPGTPRNLFGFKDGTVNPRSDAEFDAQVWNDDGGTCMIVRRVAFDMPEWES<br>VDRGTREVAMGRTIVEGAPLSGGGEFSDVDVNKLGHDGLPLIDTHSHVAMATSRNGDAERMLRRAY<br>NYDLPVTAGPAGLQDAALIDLSDTGLIFTCFQRDPDTAFIPVQRRLADGDRLNEWITHVGSAVFHVPG<br>GTTEDEYWGEALLG* |

| | | |
|---|---|---|
| *Corynebacterium variabile* | CV | MDTNFRVTRRGFLAGLGLVSATAAVGATTACAGDSDPDPVSNTDTTVPFDGPHQAGIATPLQRHNTTVAFTLRDGVDATGIRRLLRIWTGDARRLTQGEPVLADLEPELASDPGRLTVTCGFGRGLFEKARLTDKSPDWLEPLPGFAGDRLEARWGDRDLVLQICGEDRTTVSHALRVLVRGGADYARPSWTQTGFLDIPQDADGTPGTPRNLFGFKDGTVNPRTDAEFDDQVWISGEAATHPSQIGGTCMIIRRIAFDMPLWESADRPTREVSMGRTIVEGNPLSGEKEHDDPDTAAVGADGLPVIDRNAHIALAGVHDGDTRQRMLRRAYNYDLPVTPSSADALVDADPVALSDTGLIFICFQPDPRTSFIPVQRRLAAGDRLNEWITHVGSAVFLVPAGTTEEEYWGEALLG |
